# Characterization of the cryptic interspecific hybrid *Lemna × mediterranea* by an integrated approach provides new insights into duckweed diversity

**DOI:** 10.1101/2023.10.19.562947

**Authors:** Luca Braglia, Simona Ceschin, M. Adelaide Iannelli, Manuela Bog, Marco Fabriani, Giovanna Frugis, Floriana Gavazzi, Silvia Gianì, Flaminia Mariani, Maurizio Muzzi, Emanuele Pelella, Laura Morello

**Affiliations:** Institute of Agricultural Biology and Biotechnology, National Research Council, Via Bassini 15, 20133 Milan, Italy; Department of Sciences, University of Roma Tre, Viale G. Marconi 446, 00146 Rome, Italy; NBFC -National Biodiversity Future Center, 90133 Palermo, Italy; Institute of Agricultural Biology and Biotechnology, National Research Council, Via Salaria Km. 29,300, 00015 Monterotondo, Rome, Italy; Institute of Botany and Landscape Ecology, University Greifswald, Soldmannstr. 15, 17489 Greifswald, Germany

**Keywords:** Duckweed, aquatic plants, interspecific hybrids, cytotype, *Lemna gibba*, *Lemna minor*, DNA barcoding, morphometry

## Abstract

Lemnaceae taxonomy is challenged by the particular morphology of these tiny free-floating angiosperms, reduced to a single leaf-like structure called frond, without or with one to few roots. Although molecular taxonomy has helped clarify the phylogenetic history of this family, inconsistency between morphological data and nuclear and plastid markers still poses challenging questions in some cases, leading to frequent misclassifications in the genus *Lemna*. Recently, the finding that *Lemna japonica* is an interspecific hybrid between *Lemna minor* and *Lemna turionifera*, provided a clear explanation to one of such taxonomic questions. Here we demonstrated that *L. minor* is also capable to hybridize with *Lemna gibba*, generating a cryptic, previously unrecognized, but widespread taxon in the Mediterranean area. The nothotaxon *Lemna* × *mediterranea* is described through the detailed investigation of seven hybrid clones from a living germplasm collection and compared with clones of the putative parental species *L. minor* and *L. gibba*. Genetic analysis revealed that two different cytotypes, diploid and triploid, originated by at least two independent hybridization events. Despite high overall similarity, morphometrical, physiological and biochemical analyses showed an intermediate position of *L.* × *mediterranea* between its parental species in most qualitative and quantitative characters, and also separation of the two hybrid cytotypes by some criteria. These data provide evidence that hybridization and polyploidization, driving forces of terrestrial plant evolution, contribute to the duckweed genetic diversity and may have also shaped the phylogenetic history of these mainly asexual, aquatic plants. Further elucidation of hybridization mechanisms and flowering regulation will provide perspectives for future breeding strategies.

## INTRODUCTION

The Lemnaceae family is exclusively composed of aquatic plants (commonly named duckweeds) that are the smallest flowering plants, showing a body plan reduced to a single leaf-like structure called frond, without or with one or few roots. Main morphological traits are limited to frond shape, size and colour, root number and length, and position and number of vegetative pouches (Landolt, 1986). Additional diagnostic traits are vein number, the presence of a prophyllum at the base of the root(s) or papules on the dorsal side of the frond. Flowers, fruits and seeds, provide important additional taxonomic traits but are rarely or never observed in some species, as duckweeds mostly propagate asexually by forming daughter fronds from vegetative pouches on the mother frond. Key morphologic features for each species were recently updated (Bog et al., 2020a), but classification by morphology remains in some cases insufficient as not all specimens are assignable to one of the 36 recognized species with confidence. A detailed morphometric analysis has proven helpful to distinguish the American species *Lemna minuta* Kunt, invasive in Europe, from the native *Lemna minor* L. (Ceschin et al., 2016). The problem has been partially overcome with the introduction of molecular taxonomy that provided new instruments for species delimitation. Barcoding plastid markers (Les et al., 2002; Wang et al., 2010; Borisjuk et al., 2015) and nuclear sequences, as ITS, ETS (Tippery et al., 2015) as well as AFLP (Bog et al., 2015; Bog et al., 2019), mostly contributed to an almost complete phylogenetic reconstruction of the Lemnaceae family, which includes five monophyletic genera: *Lemna*, *Spirodela*, *Landoltia*, *Wolffia* and *Wolffiella* (Les and Crawford, 1999). Nevertheless, some species remain poorly delimited, particularly in the genera *Wolffia* and *Wolffiella* (Tippery et al., 2015; Bog et al., 2019). In the genus *Lemna*, inconsistency between nuclear and plastid markers impairs taking apart clones of *Lemna japonica* Landolt (Landolt, 1980), often mistaken for *Lemna minor*: plastid barcoding sequences are in fact almost identical. This issue was recently solved by using the nuclear molecular marker TBP, based on intron-length polymorphism of the β-tubulin gene family members, which provided evidence that this species is an interspecific hybrid between *L. minor* and *Lemna turionifera* (Braglia et al., 2021a). This was recently confirmed by whole genome sequencing of three different *Lemna × japonica* clones flanked by Genomic In Situ Hybridization analysis (Ernst et al., 2023). The three taxa form a species complex (an assemblage of species, which are related morphologically and phylogenetically, so that the boundaries between them are often unclear), which includes cytotypes with different ploidy levels, under detailed investigation by a multidisciplinary approach including pangenome analysis, genome size measurement, karyotype analysis in combination with physiological aspects (Abramson et al., manuscript in preparation).

*Lemna minor* also shares many morphological traits with the sister species *Lemna gibba* L. and distinction of the two may be challenging in some cases. Usually, *L. gibba* specimens are easily identified for the pronounced gibbosity of the ventral side of its fronds, due to a diffused and inflated aerenchyma, but this trait is partially influenced by growth conditions that in some cases do not make it as noticeable (Landolt, 1986). In addition, intermediate forms that cannot be determined with certainty have been reported in The Netherlands (De Lange and Pieterse, 1973; Kandeler, 1975; Landolt, De Lange and Westinga, 1979) so that the two species have been described as forming a species complex (De Lange et al., 1981). Interestingly, a new species similar to *L. gibba* was described in Italy in 1973 under the name *L. symmeter* Giuga (Giuga, 1973). However, the description of this species was not validly published following the criteria of the time (no Latin description), and almost forgotten. *Lemna symmeter* had been identified at several sites along the coast of the Campania region (Southern Italy) and described as similar to the strongly globose *L. gibba*, but only slightly ventricose and with smaller aerenchyma spaces. In particular, the two species were described as easily distinguished for the symmetric growth of the two stamens in *L. symmeter*, compared with the asynchronous growth in *L. gibba*. While *L. gibba* was reported to produce fruits and seeds, *L. symmeter* was described as sterile, producing abortive ovules and indehiscent anthers (Giuga, 1973). Kandeler (1975) hypothesized that *L. symmeter* could be an interspecific hybrid between *L. gibba* and *L. minor*, as also later reported by Landolt (1986), but this possibility was never investigated thereafter.

More recently, non-gibbous forms of *L. gibba*-like specimens of uncertain taxonomic assignment were described at some places in Central Italy (Marconi et al., 2019). However, when analysed by plastid markers, these specimens were all assigned to *L. minor*, supporting the idea of morphologic variants of this species. One of the clones isolated during that study was sent to the Landolt collection and registered as 9562; it is analysed here and designated as the hybrid type.

The existence of natural interspecific hybrids between *L. minor* and *L. gibba* was finally hypothesized, upon a large screening of clones belonging to the *Lemna* genus present in the Landolt Duckweed Collection (Braglia et al., 2021b). Similar to *L.* × *japonica*, the new hybrid taxon was first identified on a molecular basis by TBP fingerprinting and reported with the hybrid formula *L. gibba* × *L. minor*. This finding accounts for the erroneous species assignment using plastid markers of maternal origin.

The main aims of this paper are: (i) to fully demonstrate on a genetic basis the hybrid nature of the six clones previously identified, plus an additional one (LM0027) successively recovered from the Botanical Garden of Naples (Italy), and (ii) to characterize this interspecific *Lemna* hybrid based on morphological, physiological and biochemical traits in comparison with clones of the two parental species. Such characterization is supported by molecular analysis of plastid and nuclear markers of the six original clones of the Landolt Collection plus an additional one coming from the Botanical Garden of Naples (Italy).

## MATERIALS & METHODS

### Plant material

Seven putative hybrid clones, here assigned to the hybrid taxon *L.* × *mediterranea*, were analysed in comparison with several clones of the two parental species, *L. minor* and *L. gibba* by different approaches. Most analysed clones originated from the historical living plant collection of Prof. Elias Landolt (Lammler and Bogner, 2004), presently maintained as part of the IBBA collection (Milano, Italy), while others came from other collections in Europe or were collected in Italy by the Authors and integrated into the IBBA collection. All clones are listed in Table 1 with the name of the donor, collection site and date, and the experiment in which they have been used.

**Table 1.**
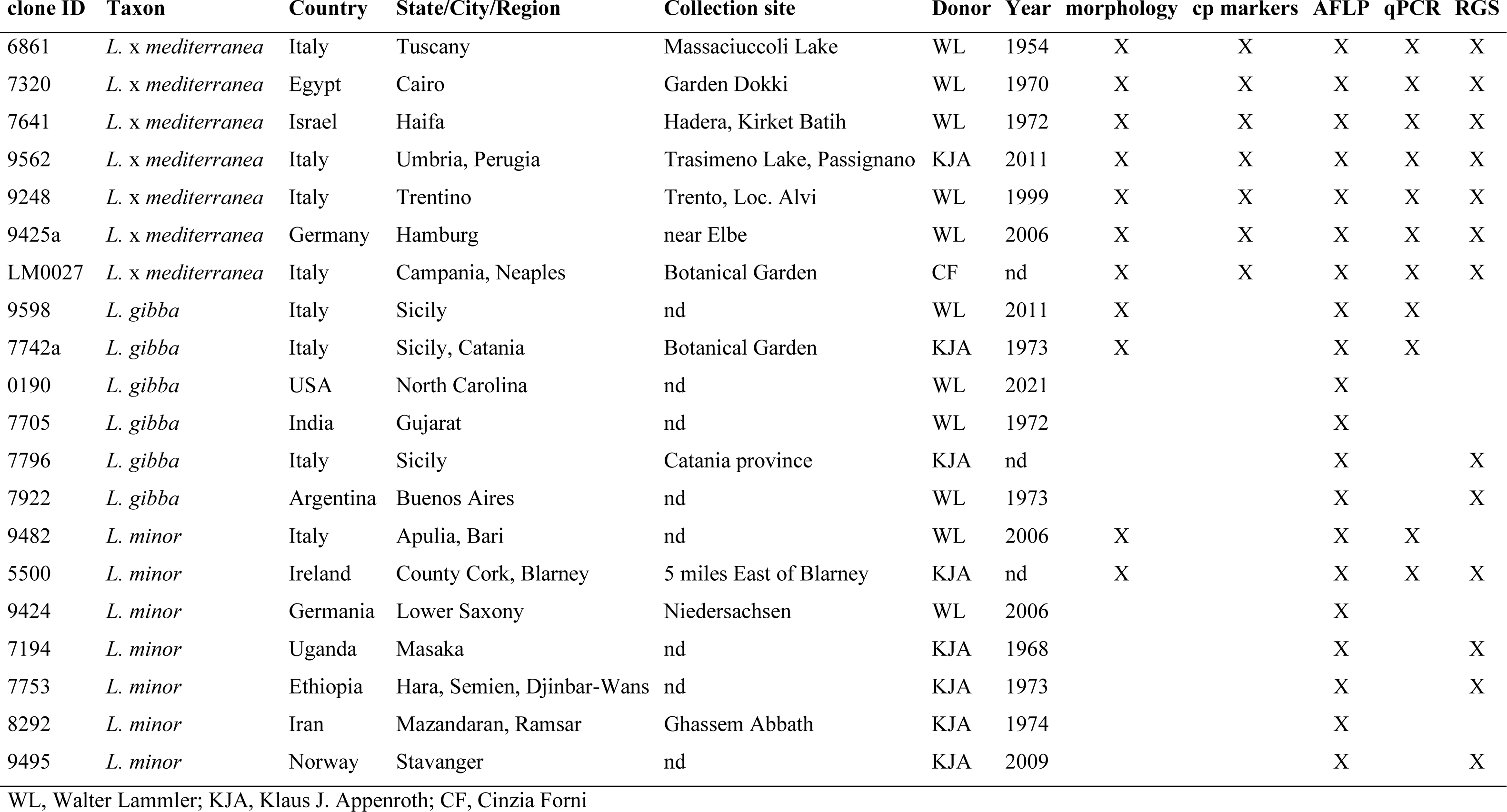
List of analysed accessions.

| clone ID | Taxon | Country | State/City/Region | Collection site | Donor | Year | morphology | cp markers | AFLP | qPCR | RGS |
| --- | --- | --- | --- | --- | --- | --- | --- | --- | --- | --- | --- |
| 6861 | *L. x mediterranea* | Italy | Tuscany | Massaciuccoli Lake | WL | 1954 | X | X | X | X | X |
| 7320 | *L. x mediterranea* | Egypt | Cairo | Garden Dokki | WL | 1970 | X | X | X | X | X |
| 7641 | *L. x mediterranea* | Israel | Haifa | Hadera, Kirket Batih | WL | 1972 | X | X | X | X | X |
| 9562 | *L. x mediterranea* | Italy | Umbria, Perugia | Trasimeno Lake, Passignano | KJA | 2011 | X | X | X | X | X |
| 9248 | *L. x mediterranea* | Italy | Trentino | Trento, Loc. Alvi | WL | 1999 | X | X | X | X | X |
| 9425a | *L. x mediterranea* | Germany | Hamburg | near Elbe | WL | 2006 | X | X | X | X | X |
| LM0027 | *L. x mediterranea* | Italy | Campania, Naples | Botanical Garden | CF | nd | X | X | X | X | X |
| 9598 | *L. gibba* | Italy | Sicily | nd | WL | 2011 | X |  | X | X |  |
| 7742a | *L. gibba* | Italy | Sicily, Catania | Botanical Garden | KJA | 1973 | X |  | X | X |  |
| 0190 | *L. gibba* | USA | North Carolina | nd | WL | 2021 |  |  | X |  |  |
| 7705 | *L. gibba* | India | Gujarat | nd | WL | 1972 |  |  | X |  |  |
| 7796 | *L. gibba* | Italy | Sicily | Catania province | KJA | nd |  |  | X |  | X |
| 7922 | *L. gibba* | Argentina | Buenos Aires | nd | WL | 1973 |  |  | X |  | X |
| 9482 | *L. minor* | Italy | Apulia, Bari | nd | WL | 2006 | X |  | X | X |  |
| 5500 | *L. minor* | Ireland | County Cork, Blarney | 5 miles East of Blarney | KJA | nd | X |  | X | X | X |
| 9424 | *L. minor* | Germany | Lower Saxony | Niedersachsen | WL | 2006 |  |  | X |  |  |
| 7194 | *L. minor* | Uganda | Masaka | nd | KJA | 1968 |  |  | X |  | X |
| 7753 | *L. minor* | Ethiopia | Hara, Semien, Djinbar-Wans | nd | KJA | 1973 |  |  | X |  | X |
| 8292 | *L. minor* | Iran | Mazandaran, Ramsar | Ghassem Abbath | KJA | 1974 |  |  | X |  |  |
| 9495 | *L. minor* | Norway | Stavanger | nd | KJA | 2009 |  |  | X |  | X |

WL, Walter Lammler; KJA, Klaus J. Appenroth; CF, Cinzia Forni

### Propagation of duckweed clones

Axenic stock cultures were maintained in Petri dishes on agarized SH medium, pH 5.1 (Schenk and Hildebrandt, plus 8 g/L Plant Agar, Duchefa) supplemented with 0.1 % sucrose, under the following growth conditions: T = 18°C; photoperiod: 16 h day, 8 h night; light intensity: 80 ±10 μmol m^-2^ s^-1^. For each set of analysis/measurements, plants were transferred into liquid medium or water, as described in the specific experimental section.

### DNA Extraction and quantification

DNA extraction was performed from about 100 mg fresh weight, using the DNeasy Plant Mini Kit (QIAGEN) as reported previously (Braglia et al., 2021a) and eluted in 150 μL of 50 mM TRIS, pH 9. When necessary, DNA was more precisely quantified through the dsDNA HS Assay Kit for Qubit fluorometer (Thermo Fisher Scientific).

### Relative Genome size measurement

Relative genome size measurements were performed using a CyFlow Space flow cytometer (Sysmex Partec GmbH, Görlitz, Germany). To extract nuclei from fresh plant tissue, about 3-4 fronds of the internal standard *Lemna aequinoctialis* Welw. (6746) and 2-3 fronds of the sample were chopped carefully in 500 µl Otto I buffer (0.1 M citric acid, 0.5% (v/v) Tween 20; Ulrich and Ulrich, 1991) with a sharp razor blade. The extract was incubated for 5 min on ice and then filtered (ca. 30 µm filter size). Subsequently, 500 µl of the staining Otto II buffer (0.4 M Na2HPO4, 4 mg/ml DAPI; Ulrich and Ulrich, 1991) were added and the sample was measured after an incubation of 5 min in darkness in the flow cytometer equipped with a 375-nm UV laser. Data collection was stopped after minimum 10,000 events and the relative genome sizes were calculated as the proportion of fluorescent intensities of the sample to the internal standard.

### TBP amplification

TBP amplification, amplicon separation by capillary electrophoresis (CE) and fragment analysis were performed as reported in Braglia et al. (2023) with minor variations. Amplification of specific β-tubulin loci (*TUBB*1 and *TUBB*2) was performed according to Braglia et al. (2021a).

### DNA barcoding analysis

**The** *atp*F-*atp*H and *psb*K-*psb*I plastid intergenic spacers were investigated as DNA barcoding regions by PCR amplification followed by Sanger sequencing as reported in Braglia et al. (2021b). Species identity was inferred from BLAST analysis against the corresponding sequences of *L. minor* (5500) and *L. gibba* (7742a) reference clones. For SNPs identification, sequences were aligned using the Vector NTI alignment tool, AlignX.

### AFLP and data analysis

The AFLP analysis was performed on all 21 duckweed clones listed in Table 1 and referring to three plant groups: *L. gibba*, *L. minor* and putative hybrids *L.* × *mediterranea*. Fifty nanograms of gDNA were analysed following the protocol of Vos et al., (1995) with modifications as described in Braglia et al., (2021b) considering a double DNA digestion (*Eco*RI and *Mse*I) and performing pre-selective and selective PCR amplification steps using the primers listed in Table S1. The Capillary Electrophoresis (CE) loading mixture and running protocol were prepared and adopted accordingly to Braglia et al., (2023). The AFLP pherogram elaboration and processing was performed by Gene Mapper Software v. 5.0 (Thermo Fisher Scientific, Germany), allowing the amplicon sizing and alleles detection. For scoring all the nine primer combinations (PCs), the RFUs peak detection threshold was fixed above 250 and a size range was considered between 70 and 450 base pairs. The peak size (base pairs) and height (RFUs) of each electropherogram were collected through a Microsoft Office Excel file and all the AFLP profiles were aligned according to the peak size. A binary matrix was then generated for each PC by the scoring for the presence/absence of homologous bands (0/1 respectively). FAMD - Fingerprint Analysis with Missing Data program, v.1.31 (Schlüter & Harris, 2006) was used to estimate genetic parameters: percentage of polymorphic markers, number of fixed markers, number of private markers found in each group, within-groups mean gene diversity (HS) and Nei’s (1973) between-groups gene diversity (G_ST_). Pearson’s correlation was calculated by Past 4 software v. 4.13 for Windows (Hammer et al., 2001) in order to estimate the linear association between the analysed clones. A principal component analysis (PCA) was also performed using the same software. A neighbour-net diagram was constructed using SplitsTree v. 4.19.0 (Huson and Bryant, 2006) applying the Nei-Li coefficients (Nei and Li, 1979). Two-thousand replicates were considered when performing the bootstrap analysis. The presence/absence matrix was also analysed by a more general Bayesian clustering approach using Structure v. 2.3.4 (Pritchard et al., 2000) and a more specific one for hybrid detection using NewHybrids v. 1.1 (Anderson and Thompson, 2002). As a first step, the initial matrix, which consisted of 1671 loci, was reduced to 694 loci by applying a minimum allele frequency of 25%, since the high proportion of loci with a low allele frequency hampered the Structure analysis to converge. The final dataset was run as diploid data with recessive alleles for the number of K clusters ranging from 1 to 5, with 50,000 burn-in steps and 50,000 additional steps. In total 10 repetitions for each K were run. The results from Structure were analysed by the Delta K method (Evanno et al., 2005) as implemented in StructureHarvester (Earl and von Holdt, 2012). Clumpp v1.1.2 (Jakobsson and Rosenberg, 2007) was used to average the 10 repetitions for each K for visualisation. For the NewHybrids analysis, five datasets were created, each with 200 randomly selected loci from the Structure dataset, as NewHybrids only runs stable for a limited number of loci. After a burn-in of 10,000 steps, additionally 20,000 steps were collected. Finally, the results of the five runs were averaged.

### Homoeolog-specific qPCR

The following procedure is an adaptation of the technique described as double-mismatch allele-specific (DMAS) qPCR for SNP genotyping (Lefevre, 2019). Instead of discriminating homo/heterozygous loci differing for one SNP, the technique is here applied to assign triploid hybrid clones to any of the two possible subgenome compositions, either two chromosome sets from *L. gibba* and one from *L. minor* (GGM) or vice versa (MMG). The assay, selectively targeting a short fragment of the *TUBB*2 locus in either *L. gibba* or *L. minor* genome, includes two slightly different primer pairs, one for each species-specific target, with similar annealing temperatures (60°C). Primer sequences are reported in Table S1. In the genome of hybrids, the two primer pairs are therefore homoeolog-specific, although amplification on the non-target homoeolog occurs at higher Cq. The principle is that, in parallel PCR amplifications, absolute ΔCq between the two primer pairs (Cq_minor_-Cq_gibba_) is maximal for both target species *L. minor* and *L. gibba*, homozygous at this locus, and close to zero for homoploid hybrids, where both subgenomes are equally present, behaving as heterozygous. Intermediate subgenome compositions in triploid hybrids should produce higher or lower ΔCq_(minor-gibba)_ values with respect to the diploid hybrids, respectively, depending on the prevalent subgenome.

PCR amplification was performed in a CFX-connect qPCR system (BIORAD) with hard-shell-96 well plates (BIORAD). Each reaction was carried out with 4 μL master mix (Titan HotTaq EvaGreen, BIOATLAS), 0.5 μL of each primer (from a 100 μM stock) and 3 μL of DNA (2 mg/mL), in a final volume of 50 μL. The two-step amplification profile used was the following: initial denaturation, 15 min at 95°C, followed by 39 cycles of 15 sec 95°C/60 sec 60°C and final denaturation by 0.5°C step-increase up to 95°C for melting curve analysis. Primers are listed in Table S1.

The threshold for Cq determination was set by the regression method. Primer specificity and amplification efficiency were first tested on serial dilutions (2, 0.2, 0.02 mg/mL) of gDNA purified from each parental species, *L. gibba* clones 7742a and 9598) and *L. minor* (clone 5500 and 9482), accurately quantified fluorometrically in duplicate, diluted to 2 mg/mL and measured again. Artificial hybrid genomes were then obtained by independently mixing gDNA from *L. minor* 5500 (M) with *L. gibba* (G) 7742a and *L. minor* 9482 (M) with *L. gibba* (G) 9598. Equimolar (1:1) DNA ratios (MG mix 1-2) mimicked homoploid hybrid genomes, while two unbalanced mixtures in 1:2 molar ratios (GGM mix 1-2 and MMG mix 1-2) simulated triploid hybrid genomes. The method was first validated by parallel PCR amplifications with the two primer pairs on the six artificial hybrid genomes. For statistical significance, ΔCq of each group (MG, MMG and GGM) were averaged and analysed by one-tailed ANOVA. The DNA of the two target species and the seven hybrid clones was then tested in triplicate in at least two independent experiments, by the same parallel PCR amplification. For each sample, ΔCq of all 9 replicates, excluding outliers (±2Cq from the mean) were mediated and plotted. The difference of the Cq means between triploid and diploid *Lemna* clones was tested by Student’s T testing and ANOVA.

### Morphological analyses

Morphological analyses were carried out on fronds of each of the seven putative hybrid *Lemna* clones assigned to the hybrid taxon *Lemna* × *mediterranea*, that were grown in the laboratory for three weeks, under uncontrolled temperature and light conditions, in 600 ml glass beakers filled with mineral water of known composition (Table S2). For comparison, two diploid clones of each parental species *L. gibba* (clone 7742a and 9598) and *L. minor* (clone 5500 and 9482), of European origin, were similarly grown and analysed. The entire set of beakers was placed near the window to be exposed directly to natural light respecting the summer seasonal photoperiod.

To morphologically describe the putative hybrid clones, 10 specimens of each clone were randomly collected in parallel with those of the parental species, for a total of 110 fronds. Each of these specimens were observed and photographed in dorsal, ventral and lateral position under a stereomicroscope (Olympus SZX2-ILLT) equipped with an Olympus OM-D EM-5 camera. Morphological traits of each specimen were analysed and measured using the image-processing program ImageJ software v. 1.53t (Schneider et al., 2012). The analysed traits were selected after consulting reference literature related to *Lemna* species (e.g., Landolt, 1980, 1986; Ceschin et al., 2016; Bog et al., 2020b). They included both quantitative and qualitative morphological characters, as listed below (Fig. 1). If the specimen consisted of contiguous fronds (colony), the traits were analysed only on the mother frond; it was complete with root, and was the largest and placed above all the other fronds.

**Figure 1.**
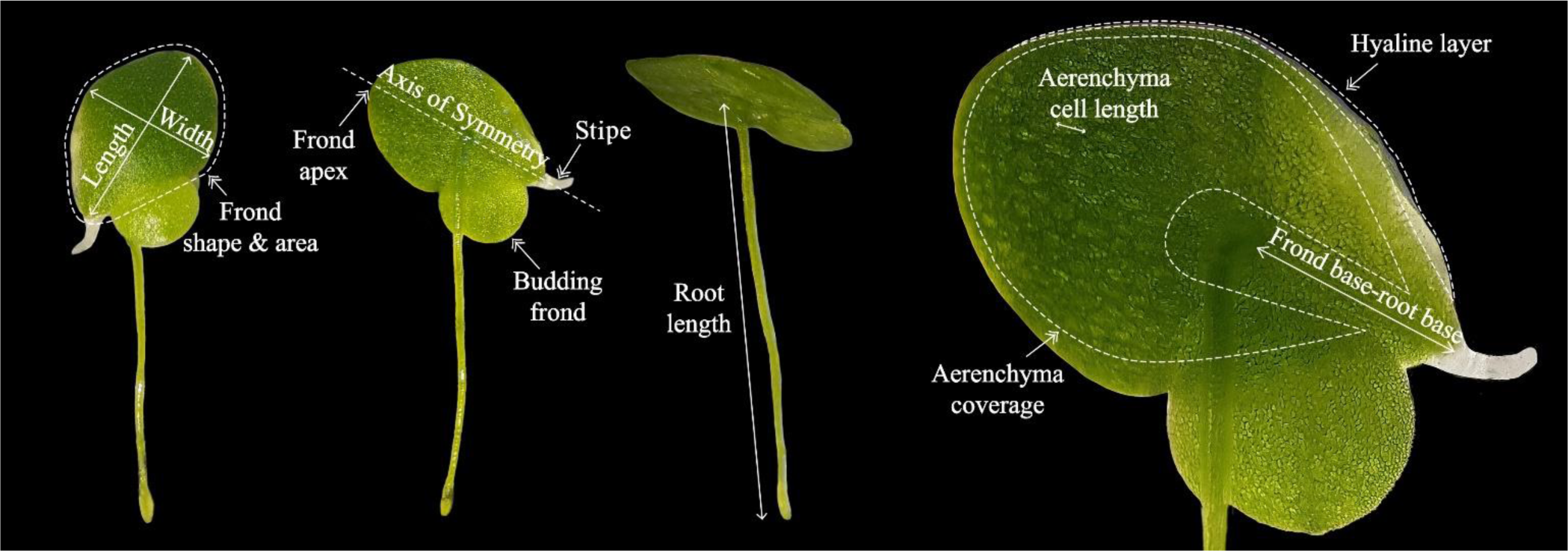
Illustration of the morphological traits analysed.

#### Quantitative traits

- frond length and width (mm)

- frond length/width ratio

- frond area (mm^2^)

- vein number (n)

- root length (mm)

- distance frond base-root base (mm)

- aerenchyma percentage coverage in frond (%)

- aerenchymatic cell length (mm)

#### Qualitative traits

- frond shape (obovate/pear-shaped/bilobate irregular/rhomboid/stocky rhomboid)

- frond symmetry (symmetrical/asymmetrical)

- frond apex (rounded/acuminate)

- hyaline layer on the frond edge (no hyaline layer/basal hyaline/central-basal hyaline/all around hyaline)

- aerenchyma position related to frond area (dispersed/central/upper-central)

- aerenchyma reaching or not the frond edge

- papules (absent/unclear/evident)

- connection stipe (presence/absence)

### Scanning electron microscope (SEM) observations

From each of the four *Lemna* groups identified by genome size measurement, 10 specimens were randomly taken and fixed overnight at 4°C in a mixture of 2% paraformaldehyde and 3% glutaraldehyde in 0.1 M cacodylate buffer. The next day, specimens were thoroughly washed in the same buffer and post fixed in 1% buffered osmium tetroxide for 90 minutes at 4°C. After thorough washing, first in 0.1 M cacodylate buffer and then in double-distilled water, the specimens were dehydrated through a graded ethanol series (15, 30, 50, 75, 85, 95 and 100%) and dried in a Critical Point Dryer (CPD 030 unit, BalTec, Balzers, Liechtenstein). Specimens were mounted on aluminium stubs using double-sided carbon discs and gold sputtered using a K550 sputter coater (Emithech, Kent, UK). The specimens were then observed and microphotographed by scanning electron microscope (SEM) (Gemini 300, Carl Zeiss AG, Jena, Germany).

### Analysis of plant growth and biochemical parameters

Plant growth and biochemical analyses were performed on cultures grown under controlled and axenic conditions in 150 x 75 mm (d x h) Petri dishes (Corning Inc., Corning, NY, USA) that contained 150 mL of freshly prepared, liquid SH medium (pH adjusted to 5.5) and 0.5% sucrose. Plants were cultivated at a 16 h day photoperiod under 100 μmol photons m^-2^ s^-1^ at 25 °C ± 2 °C. Experimental cultures were started by inoculating 30 colonies with 2-3 fronds each. Growth measurements and biochemical analyses were carried out after seven days. All cultures were set up in quintuplets.

### Frond vascular organization

To determine frond vein numbers, ten duckweed colonies, with two/three fronds each, were washed with deionized water and cleared with 70% ethanol for three weeks prior to observations using a Nikon stereomicroscope (Nikon SMZ1000) equipped with a Nikon digital camera (DS-5M; www.nikoninstruments.com/). Duckweed colonies were observed under bright and dark field conditions at 20x and 10x magnification.

### Stomatal traits

To characterize stomatal traits, for each *Lemna* clone, three colonies with 2 or 3 fronds were washed in demineralised water and immersed in 70% ethanol solution for three weeks to remove any pigmentation. Stomata features were examined and photographed using a Nikon microscope (ECLIPSE 80i) equipped with a digital camera (Nikon DS-5M; www.nikoninstruments.com/). Stomatal density and stomata size were determined by analysing images of four different microscopic fields (0.95 μm^2^) for each mother frond of three colonies/clone at a magnification of 20x. Fields were selected in the regions located between the main vein and the closest secondary vein (two sectors to each side of the main vein) (Fig. S1). Stomatal density (SD) was expressed as stomata number/area of one microscopic field (area). The size of stomata was measured using the LeafNet software (Li et al., 2022). Parameters were adjusted by setting “Stained Denoiser” for the Image denoiser function and “StomaNet Universal” for the Stoma detector function. Stoma minimum size was set to 300 μm^2^.

### Analyses of growth parameters

For fresh and dry weight measurements, all plants from each tested clone were sieved out of the medium, dry blotted and either weighed immediately (fresh weight) or dehydrated at 60 °C for 72 hours and then weighed (dry weight).

The mean single frond fresh weight of each clone was estimated by measuring the total biomass of each experimental culture and dividing by the corresponding total number of fronds (including daughter fronds when still attached to the mother) previously counted using the ImageJ image processing program (Schneider et al., 2012).

### Relative Growth Rate

The relative growth rate (RGR) of each *Lemna* clones was measured after seven days and calculated using the following formula: RGR = ln (DW_f_-DW_i_)/T_f_ -T_i_ where: DW_f_ = final dry weight (g), DW_i_ = initial dry weight (g), T_f_ = total incubation period (day), T_i_ = initial time (day). The results were expressed as g g^-1^ day^–1^.

### Determination of chlorophyll and carotenoid contents

Fresh fronds (0.1 g) were grounded into powder with liquid nitrogen, and then homogenized with 80% (w/v) cold acetone, centrifuged at 5000x*g* for 10 min. The absorbance of the supernatant was measured at 663, 646 and 470 nm. Chlorophyll a, b and carotenoids content were determined using the equations described in Lichtenthaler (1987). The results were expressed in mg of chlorophyll or carotenoids per gram of plant tissue fresh weight (mg/g FW).

### Measurement of protein content

*Lemna* fronds (0.1 g fresh weight) were grinded in liquid nitrogen with mortar and pestle. The proteins were then extracted at 4 °C with a cold 0.5 M potassium phosphate (pH 7.0) buffer containing 0.1% ascorbic acid, 0.1% Triton X-100, 1 mM EDTA and 7.5% polyvinylpyrrolidone. The homogenate was centrifuged at 4 °C for 20 min at 12000×*g*. The total soluble proteins were quantified according to Bradford (1976) using albumin bovine serum as standard. The results were expressed in mg of proteins per gram of fresh weight plant tissue (mg/g FW).

### Statistical analyses (for morphological data)

All statistical analyses were performed using R software, vers. 4.2.1. (R Core Team, 2022). All selected morphological traits and datasets comprising growth analysis and biochemical parameters were analysed using Principal Component Analysis (PCA), and biplots were made considering PC1 and PC2 using either ggfortify or the factoextra packages of the R software (Tang et al., 2016; Kassambara and Mundt, 2020). For plant growth and biochemical data analyses, the cos2 values were considered. A high cos2 value indicates a higher impact of the Wtraits were compared between clones using ANOVA. Where assumptions of normality and homoscedasticity were not met, a non-parametric test was conducted (Kruskal-Wallis). Qualitative traits were analysed by calculating contingency tables and performing Pearson’s Chi-squared tests. Boxplots and mosaicplots were made using ggplot2 package v. 3.4.2 (Kassambara, 2023). Specifically, for multivariate analyses “ade4” and “vegan R packages were used (Dray & Dufour, 2007; Oksanen et al., 2020) and the significance level was set to P < 0.05. The post hoc Tukey’s Honest Significant Difference test (TukeyHSD) was run to adjust P-values for multiple comparisons to determine which samples have significantly different means in paired sample comparison.

## RESULTS

### Molecular characterization of the additional, putative hybrid, *Lemna* clone LM0027

The hybrid origin of six of the seven clones analysed in this study from an interspecific cross between *L. minor* and *L. gibba* was previously suggested, relying on TBP profiling and plastid marker sequences (Braglia et al., 2021b). The six specimens were all identified as *L. gibba* by their collector E. Landolt, by morphologic analysis. An additional clone included in this study, LM0027 was instead classified as *L. minor* by its collector (C. Forni, personal communication to M.A.I.). However, the same TBP pattern as that observed for the other six hybrid clones, which merges profiles of the two putative parents *L. minor* and *L. gibba*, was observed for LM0027 (not shown). Every putative hybrid clone is then heterozygous at all six β-tubulin loci (Braglia et al., 2021b). LM0027 groups together with the other six putative hybrid clones by cluster analysis of TBP markers, well separated from the clusters of each parental species (seven clones each, from different geographic areas were chosen as representative of the intraspecific genetic diversity, Fig S2). Sequences of both intronic regions of the β-tubulin locus *TUBB*1 (Supplementary File 1), amplified by specific primers, confirmed also for clone LM0027 the identity of each homoeologous alleles with the corresponding parental species, upon BLAST DNA analysis against the genome sequence of *L. gibba* 7742a and *L. minor* 9252, respectively (www.lemna.org/blast; accessed on 04/27/2023). BLAST DNA analysis of the nucleotide sequences obtained for the two plastid markers *psb*K-*psb*I (512 bp) and *atp*F-*atp*H (529 bp) (Supplementary File 1) permitted to establish the parentage of the newly investigated clone LM0027, which turned out to have plastid marker sequences matching those of *L. minor*, and are almost identical to the four previously analysed hybrid clones 7641, 6861, 9562, 7320 (1 SNP), thus having *L. minor* as the maternal parent. For the two remaining clones, 9248 and 9245a, their origin from the reciprocal cross was previously assumed from their plastid marker identity to *L. gibba* sequences (Braglia et al., 2021b).

### Genome size estimation and subgenome composition of hybrid clones

Plant interspecific hybrids are in most cases polyploid but can be also diploid (homoploid) when the two different subgenomes are shared within the same nucleus without chromosome number increase (Abbott et al., 2010). The Relative Genome Size (RGS) of each *L. × mediterranea* clone was then assessed by flow cytometry in comparison with that of the parental species and used as a proxy of ploidy (Table 2). The five clones with *L. minor* as the maternal parent showed an average RGS of 0.54, exactly intermediate between the values of the two diploid parental species (0.46 *L. minor*, 0.64 *L. gibba*), perfectly fitting what expected for a homoploid hybrid. Conversely, the RGS of the two clones having *L. gibba* as the maternal parent, 0.84, was about 1.5x larger, suggesting a triploid state. This led us to conclude that the analysed clones belong to two different cytotypes, most likely a homoploid and a triploid one, respectively. Both kinds of hybrids, although rarer than tetraploids or hexaploids, may occur in plants and are generally considered as bridges toward higher ploidy levels, eventually leading to hybrid speciation (Bretagnolle 1995; Tayalè and Parisod, 2013; Mason and Pires, 2015).

**Table 2.** Genetic structure of seven *L.* × *mediterranea* clones (hybrids) and parental species. G and M refer to *L. gibba* and *L. minor* subgenomes, respectively. The estimated Relative Genome Size (RGS) and the deduced ploidy are reported.

| ID | Taxon by TBP | plastid donor | RGS | ploidy | sub genome composition |
| --- | --- | --- | --- | --- | --- |
| 7796 | *L. gibba* | *L. gibba* | 0.650 | 2n | GG |
| 7922 | *L. gibba* | *L. gibba* | 0.620 | 2n | GG |
| 5500 | *L. minor* | *L. minor* | 0.460 | 2n | MM |
| 7194 | *L. minor* | *L. minor* | 0.450 | 2n | MM |
| 9495 | *L. minor* | *L. minor* | 0.460 | 2n | MM |
| 7753 | *L. minor* | *L. minor* | 0.460 | 2n | MM |
| 6861 | *L. x mediterranea* | *L. minor* | 0.540 | 2n* | MG |
| 9562 | *L. x mediterranea* | *L. minor* | 0.540 | 2n* | MG |
| LM0027 | *L. x mediterranea* | *L. minor* | 0.541 | 2n* | MG |
| 7320 | *L. x mediterranea* | *L. minor* | 0.539 | 2n* | MG |
| 7641 | *L. x mediterranea* | *L. minor* | 0.538 | 2n* | MG |
| 9248 | *L. x mediterranea* | *L. gibba* | 0.842 | 3n* | GGM |
| 9425a | *L. x mediterranea* | *L. gibba* | 0.839 | 3n* | GGM |

\* deduced from genome size

Triploid hybrids may have two different subgenome compositions: MMG or GGM, depending on the donor of the diploid gametes. Further analysis was then conducted in order to determine the subgenome composition of each hybrid clone, by a modification of the DMAS qPCR technique (Lefever et al., 2019). Genomic DNA of two clones for each parental species, the seven hybrid clones and six artificial hybrid genomes (MG 1 and 2, GGM 1 and 2 and MMG 1 and 2) obtained by mixing genomic DNA of *L. gibba* and *L. minor* in different proportions, was amplified in parallel with two homeolog-specific primer pairs and the ΔCq_(minor-gibba)_ values were recorded. Specificity of the two primer pairs is shown in Fig. 2 A where representative amplification plots are shown. Mean ΔCq_(minor-gibba)_ for *L. gibba* 7742a and *L. minor* 5500 were equal to 10.84 and ‒ 12.83, respectively (mean of triplicate technical repetitions, 3 independent experiments). Small but significant differences in ΔCq values were observed between artificial hybrid genomes made up by three different gDNA ratios, for test validation. As expected, DNA mix MG1 and MG2, with a 1:1 composition of *L. minor* and *L. gibba* DNA, showed very low ΔCq values (Mean = ‒ 0.47 ± 0.05). Both unbalanced 1:2 DNA mixtures GGM and MMG, gave mean ΔCq values significantly (p <0.01 by ANOVA calculation and Tukey HSD) higher (0.42 ± 0.04) or lower (‒1.25 ± 0.11) than MG, respectively, with a ΔΔCq between the triploid-like DNA mix and the diploid-like mix of + 0.89 and ‒ 0.78, respectively, a difference sufficient to discriminate between the two genotypes. Mean ΔCq obtained for each natural hybrid DNA were then plotted (Fig. 2 B). Mean ΔCq values obtained for the triploid group (0.75 ± 0.21 and 0.52 ± 0.32) were significantly higher (ANOVA, p < 0.05) than the mean ΔCq of the diploid group, close to 0 (‒ 0.080 ± 0.14), giving ΔΔCq values of 0.83 and 0.59 respectively, which indicates that triploids have a GGM genotype. This conclusion is also supported by the observation that the measured RGS for these two triploid clones, 0.84 (Table 2) is closer to the theoretical genome size calculation for GGM hybrids (0.86), than for MMG (0.77), based on RGS of the parental species. The small discordance between ΔCq values of hybrid clones and the corresponding artificial genome mixtures is likely due to inaccuracy of quantification of the DNA preparation used to make admixtures.

**Figure 2.**
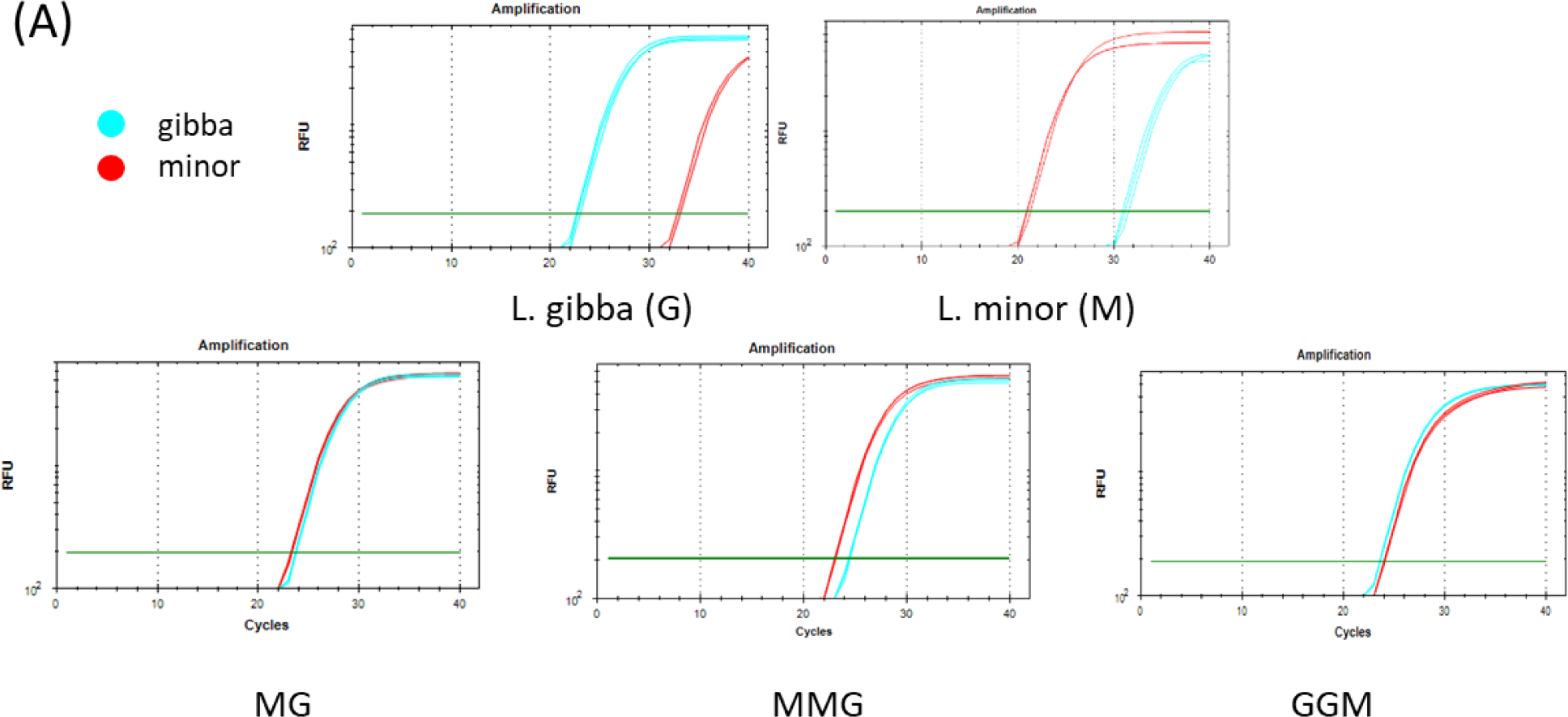
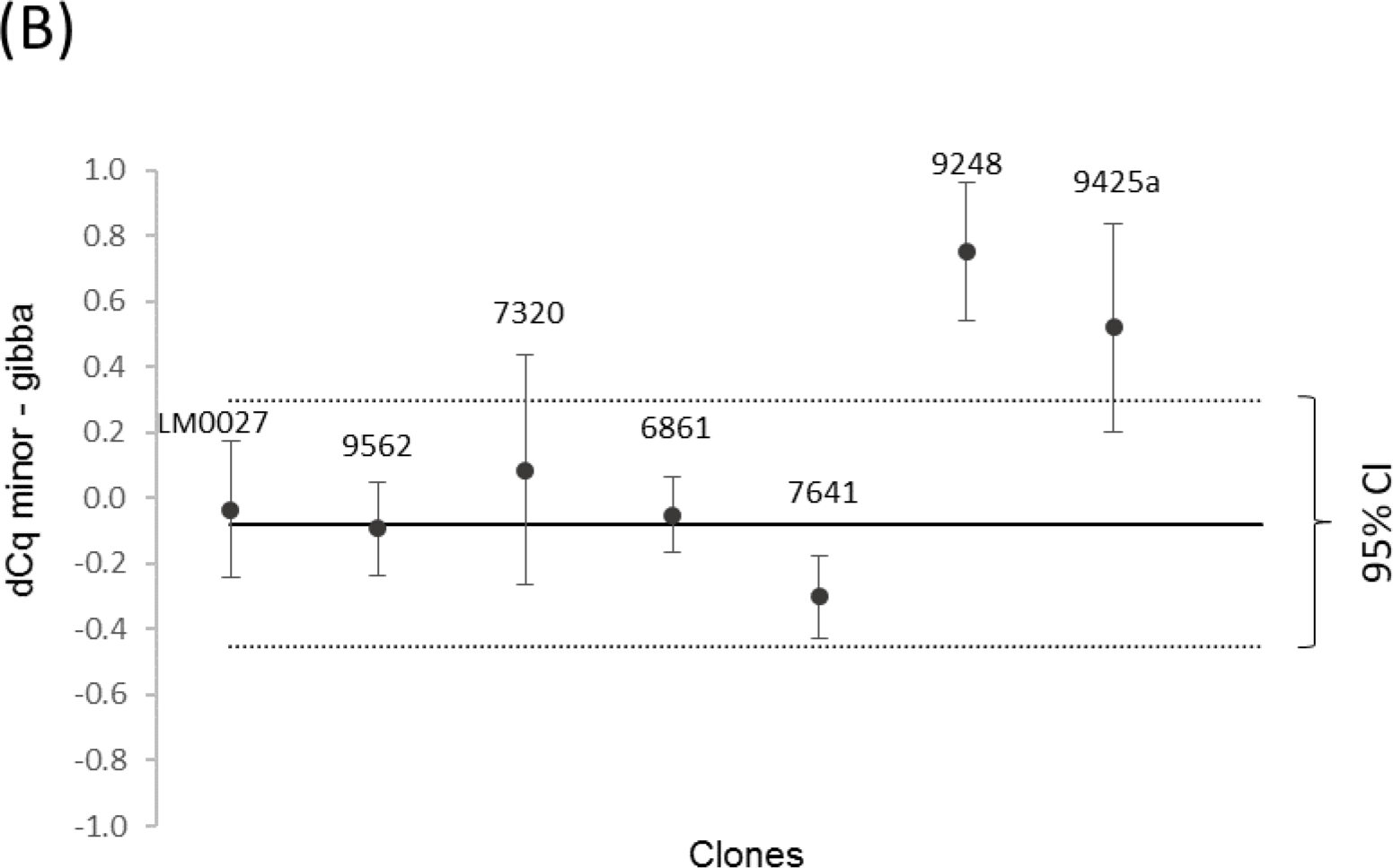
Homeolog-specific qPCR. (A) Representative PCR amplification plots of the parental species DNAs and their mixtures in different proportions (upper panel). Colours indicate the specific target of the primer pair used. (B) Scatter plot of the Cq differences between the perfect match and mismatch reactions for each *L.* × *mediterranea* clone (n=3). Horizontal lines indicate the mean value of the five diploid samples and the 95% Confidence Interval (± 2SD).

### Genome diversity by AFLP analysis

AFLP analysis of seven clones for each group (parents and putative hybrids) already analysed by TBP provided confirmation of hybridization at the whole genome scale. In this regard, the AFLP analysis yielded 1671 markers, 98% of which were polymorphic considering 21 duckweed clones. The number of polymorphic markers within the groups of clones of *L. minor* and *L. gibba* was 896 (54%) and 856 (51%) respectively, significantly higher than those estimated within the third group of clones (*Lemna* × *mediterranea*) that revealed only 21% of polymorphism. Accordingly, the lowest number of private markers, 37, was found in this latter group, compared to 456 and 354 private markers detected for *L. minor* and *L. gibba* groups respectively, reflecting the conspicuous number of loci shared between the putative hybrid group and both parents. In addition, mean genetic diversity estimated within taxa (HS) was 0.1059 in *L. gibba*, 0.0750 in *L. minor* and 0.0221 in *L.* × *mediterranea*. Conversely the Nei’s (1973) between-population gene diversity (G_ST_) value was significantly higher (p < 0.05) comparing each other the *L. minor* and *L. gibba* groups (0.2638), than comparing *L.* × *mediterranea* to either of the two parents (0.1224 and 0.1160 to *L. minor* and *L. gibba*, respectively). In this context, the diagram of the Pearson’s linear correlation (Fig. 3) estimated among all analysed clones returned the highest significantly (p < 0.05) recorded values among the accessions of *L.* × *mediterranea*, forming a group of clones strongly related to each other, while the lowest correlation was assessed among *L. gibba* clones.

**Figure 3.**
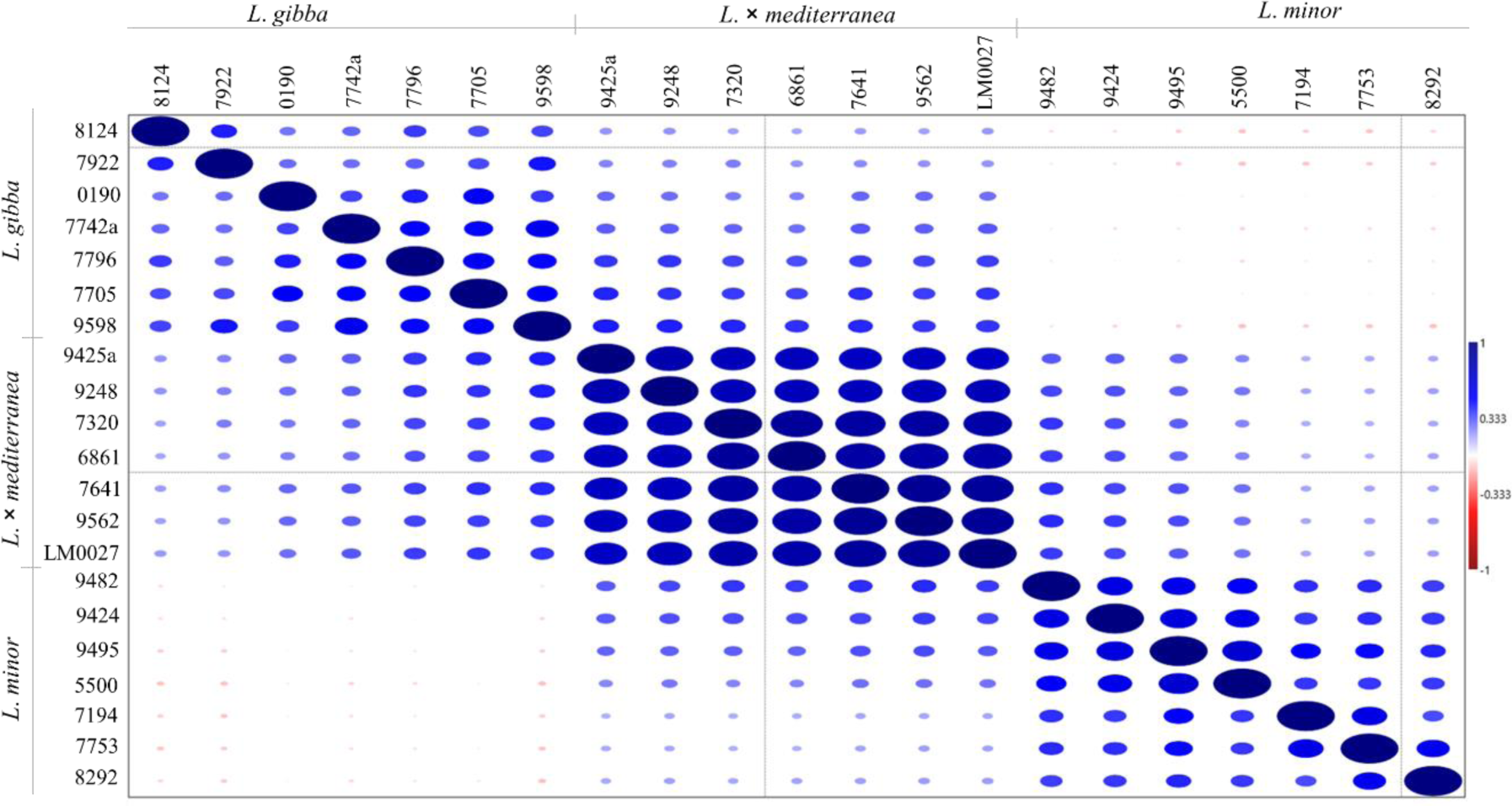
Pairwise Pearson’s correlation matrix, comparing parental and putative hybrid genotypes. Clone numbers refer to those reported in Table 1.

The total variance accounted for by each component of the PCA (PC1 37% - PC2 21%) in Fig. S3 grouped the analysed clones in three distinct, non-overlapping and well-defined clusters, further highlighting that a representative quote of the total genetic variability (52%) can be attributed to variability detected among groups.

Neighbour-net analysis (Fig. 4) also supported the existence of differentiated groups of individuals. Despite the evident reticulation, three diverging groups were formed by a strongly supported split (bootstrap values: 87, 90 and 100%): two of these correspond to the *L. minor* and *L. gibba* groups of clones, considered as the parental species involved in the cross, whereas a third group, located between the other two, represents an isolated entity formed by the seven clones of *L.* × *mediterranea* taken into account as the derived hybrid. A substantial amount of reticulation particularly occurred within parental groups, reflecting the geographic partition (America, Europe, India and Africa) (Table 1) characterizing the selected clones, accounted as representative of the two species. Moreover, within the *L.* × *mediterranea* group two sub-branches were observed, in accordance with the already documented different chloroplast origin (coloured lines in Fig. 4) and subgenome composition (GGM, triploid) of two of the seven hybrid clones (9248 and 9425a).

**Figure 4.**
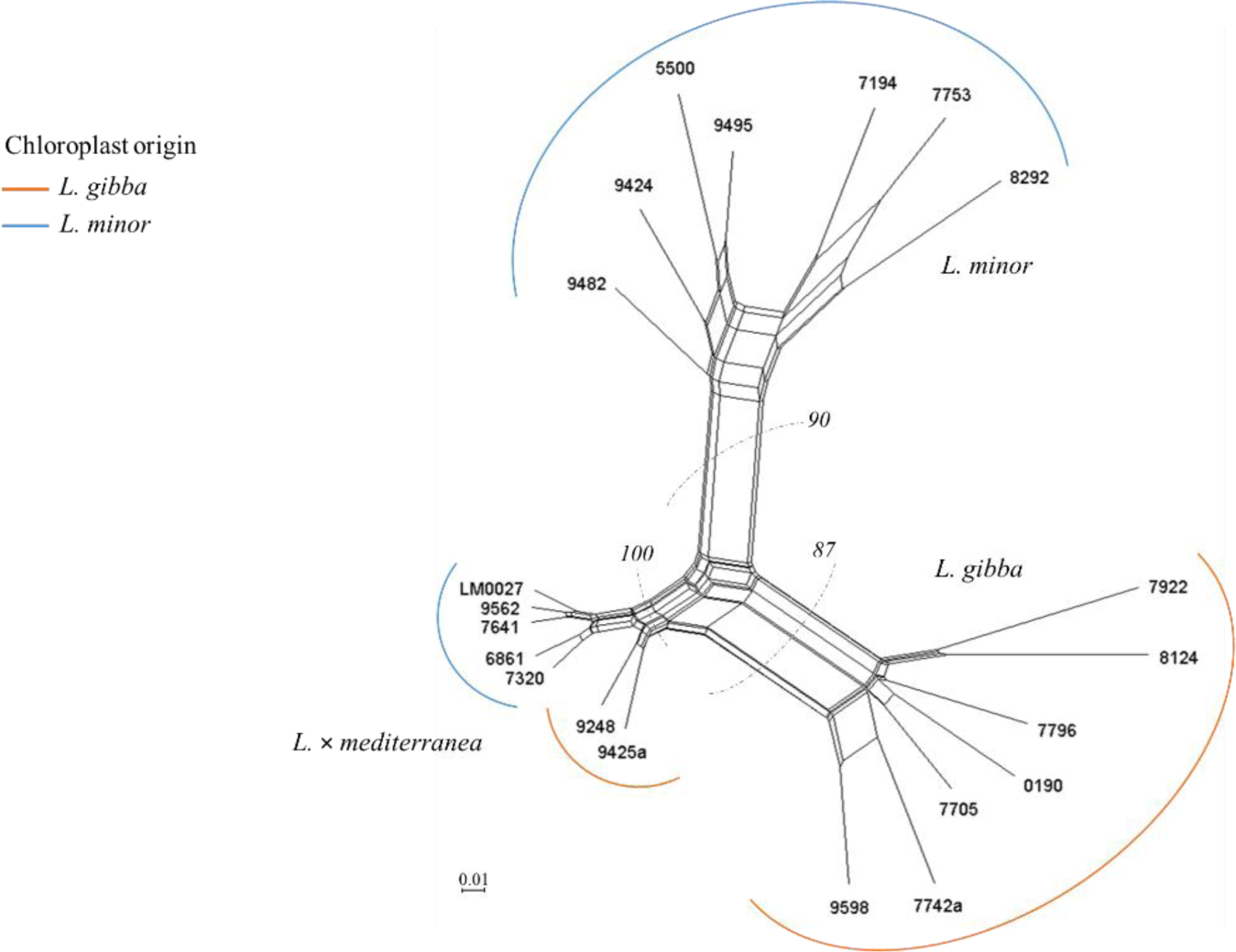
Phylogenetic network (NeighborNet) constructed on the AFLP patterns. Bootstrap values are given for the main clusters. Grouping by colours is made according to the maternal parentage determined by plastid markers.

The Structure analysis with the reduced dataset (694 loci) reinforces the origin of the putative hybrid individuals. According to the delta K method, the highest probability of dividing the data set into two clusters correlates with the two parent species. The putative hybrid individuals show an approx. 50% affiliation to each of the two clusters of the parent species (Fig. 5) and they would even be assigned to their own cluster if three clusters were assumed. Strikingly, the two individuals with GGM genome composition show a further deviation from the hybrid cluster when assuming 4 clusters.

**Figure 5.**
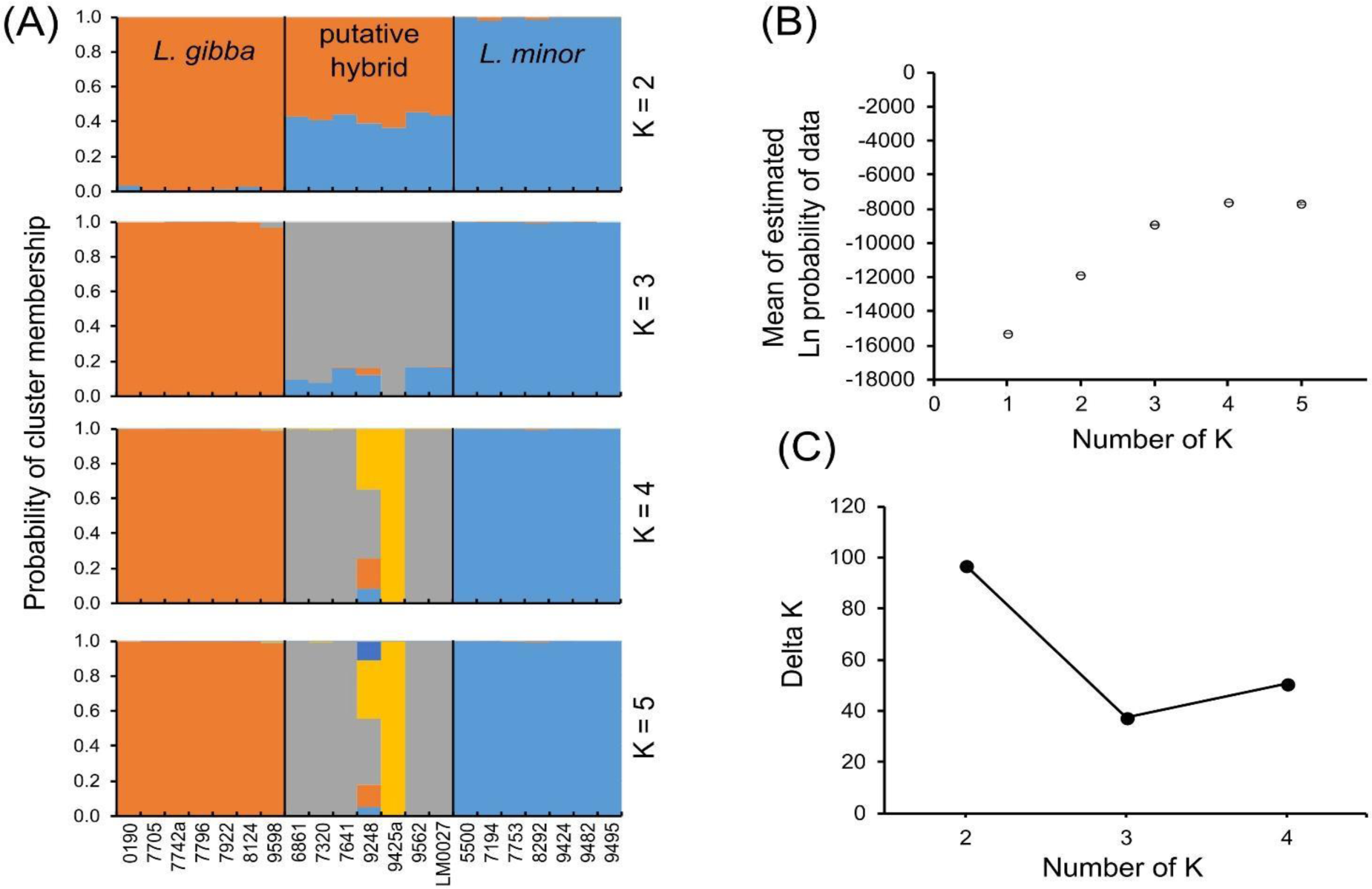
Results of the Structure analysis based on the reduced AFLP dataset. (A) Cluster membership of the 21 investigated clones for the number of clusters K=2-5. (B) Mean Ln probability values and their standard deviations from the 10 independent Structure runs for K=1-5. (C) Results of the Delta K method, showing the highest value for K=2.

These results are further supported by the analysis with NewHybrids (Table S3). All putative hybrid individuals were categorized as F1 hybrid crosses between *L. gibba* and *L. minor*. There was no assignment to the F2 or either backcross category.

### Morphological diversity

To compare morphological diversity between *L*. × *mediterranea* and the parental species, two diploid clones of Mediterranean origin, more closely related to the hybrids, were chosen as representative of each parental species *L. gibba* (GG) and *L. minor* (MM). Morphological analysis of 10 fronds of each clone of the two parents, as well as the two hybrid cytotypes, showed that the two hybrid classes, triploid (GGM) and homoploid (MG), are distinct not only genetically but also morphologically, despite large trait overlaps with one other and with parental species (Fig. 6A).

**Figure 6.**
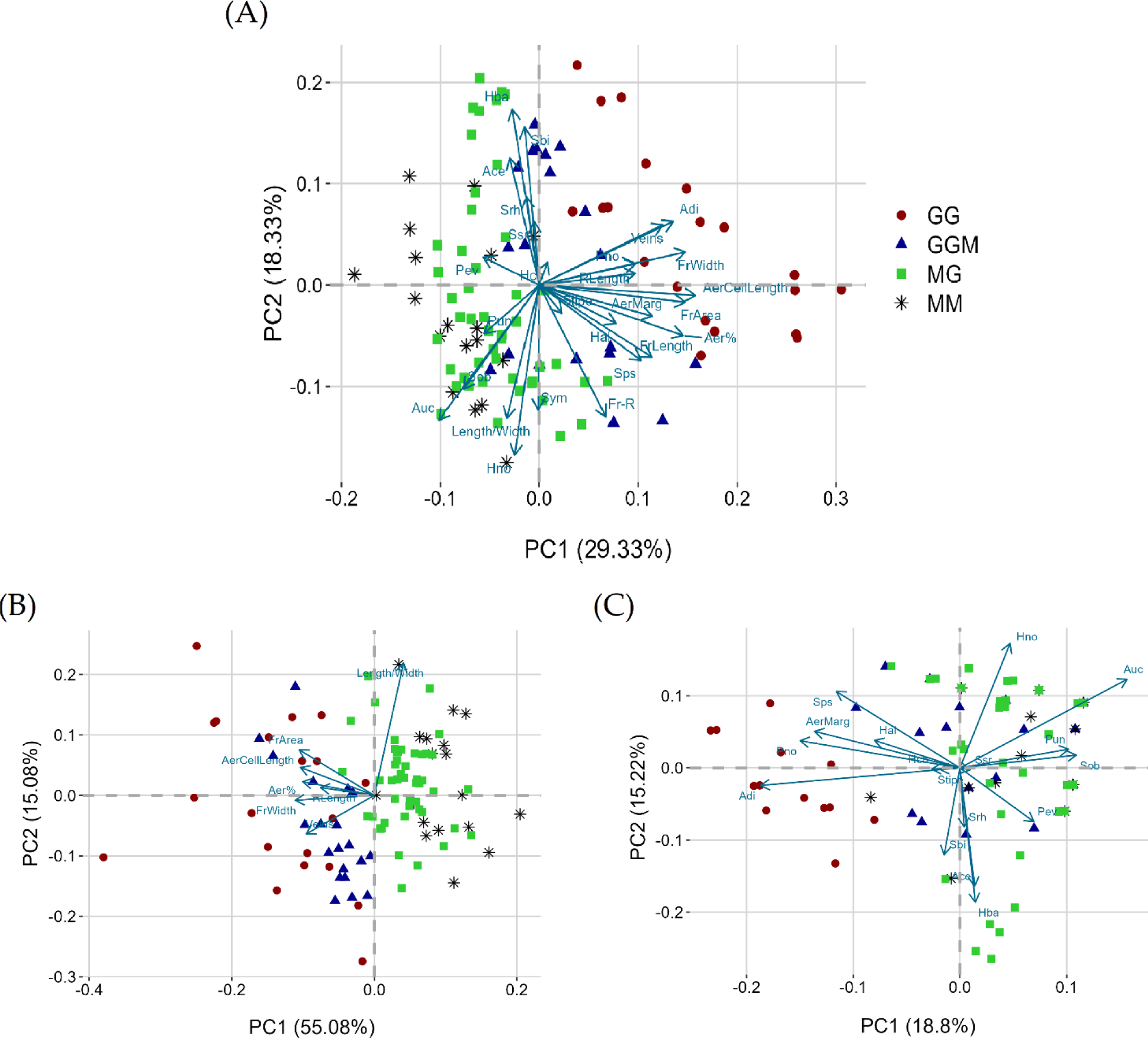
PCA based on all considered morphological traits for the 11 investigated clones (10 fronds each) (A). Quantitative (B) and qualitative (C) morphological traits statistically significant from ANOVA and Chi-squared tests, respectively, are shown. Acronyms for quantitative traits: FrLenght = frond length, FrWidth = frond width, Length/Width = frond length/width ratio, FrArea = frond area, Veins = vein number, RLenght = root length, Fr-R = distance frond base-root base, Aer% = aerenchyma percentage coverage in frond, ArCellLenght = aerenchymatic cell length. For qualitative traits: Sbi = bilobate irregular frond shape, Sob = obovate shape, Sps = pear-shaped, Srh = rhomboid shape, Ssr = stocky rhomboid shape; Hal = all around hyaline frond edge, Hba = basal hyaline edge, Hcb = central-basal hyaline frond edge, Hno = no hyaline hyaline edge, Ace = central aerenchyma position, Adi = dispersed aerenchyma, Auc = upper-central aerenchyma, AerMarg = aerenchyma reaching the frond edge, Pno = absent papules, Pun = unclear papules, Pev = evident papules.

Most of the morphological traits considered are useful in differentiating between the four *Lemna* groups (Table S4). Such differences are more marked between the two parental species than between them and the two hybrid cytotypes. There are significant differences between the two hybrid cytotypes particularly in quantitative traits (Fig. 6B), while for qualitative ones, there are several overlaps (Fig. 6C; Table S4).

All the quantitative morphological traits considered, except aerenchymal cell length, showed significant differences between the two hybrid cytotypes (Fig. 7, Table S5). GGM fronds differed significantly from MG for: larger surface (7.90 vs 6.53 mm^2^), higher width (2.87 vs 2.54 mm), higher aerenchyma abundance (72.50 vs 54.60 %), longer roots (5.87 vs 2.81 mm) and a higher number of veins (5.00 vs 3.8), on average. Conversely, GGM and MG did not show any significant difference from the maternal species *L. gibba* and *L. minor*, respectively, in relation to some quantitative parameters (frond area, frond length/width ratio, root length); in addition, GGM did not differ significantly from *L. gibba* for frond width (Fig. 7, Table S5).

**Figure 7.**
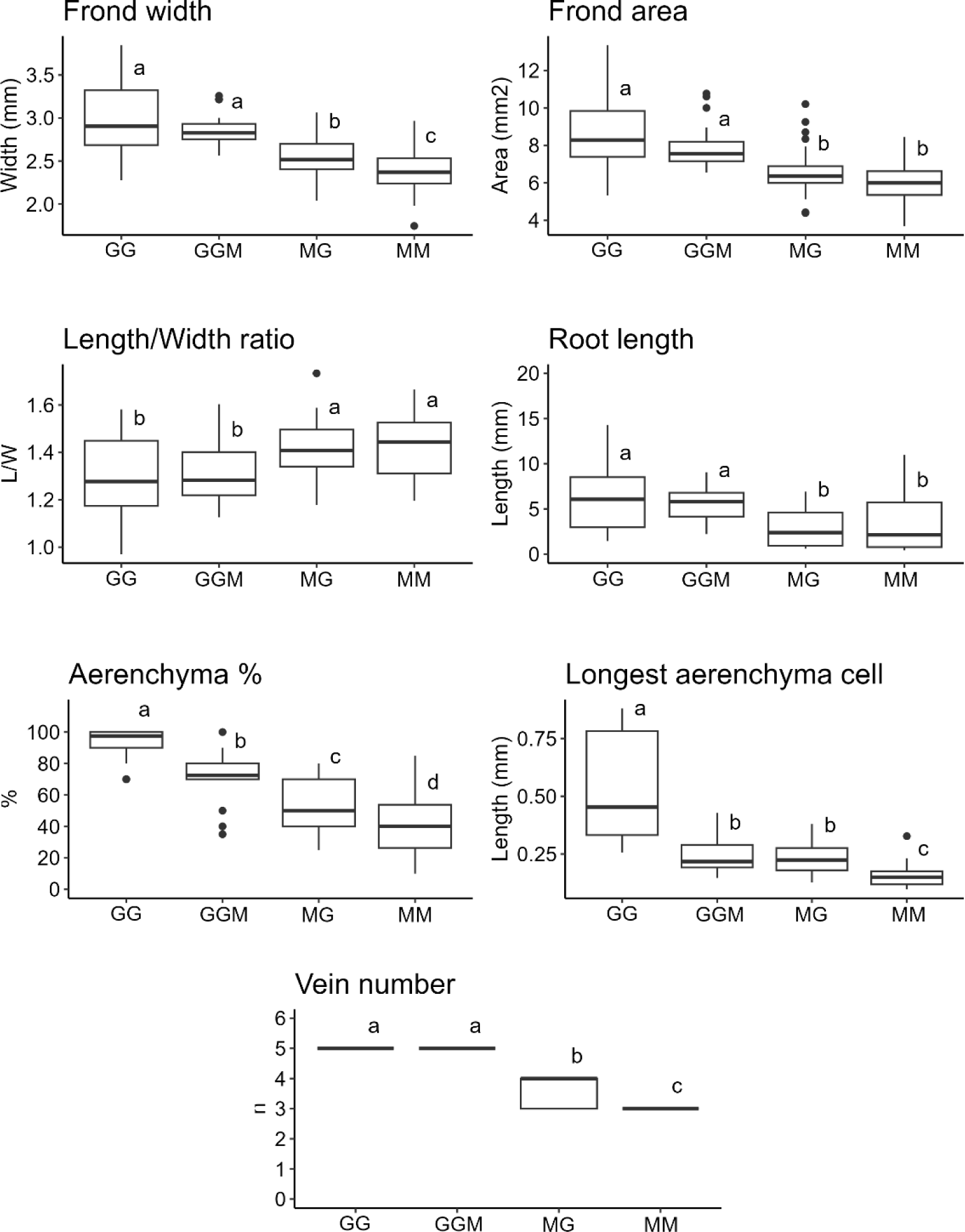
Differences among the four *Lemna* groups in relation to quantitative traits found to be significant by ANOVA test. In each graph, boxplots with different letters represent significant differences at p value ≤ 0.05.

Although there were significant differences in qualitative morphological traits between the four genetically distinct groups of *Lemna* clones studied, several overlaps for these traits were found between the two hybrid cytotypes (Fig. 8). Thus, this set of traits contributes less to differentiating the two hybrid cytotypes. With specific reference to the frond shape, the pyriform shape occurred in all the groups except for *L. minor,* which had a predominantly obovate shape; a stocky rhomboid shape was absent only in *L. gibba*. A frond edge completely hyaline all round was characteristic of *L. gibba* and GGM while it was very sporadic in *L. minor* and MG. A total absence of the hyaline edge was mainly found in both *L. minor* and MG, while it was sporadic in *L. gibba*. Elongated stipes, stolon-like appendage connection daughter and mother fronds, occurred in all *Lemna* groups, except for *L. minor*. Only in *L. gibba*, aerenchyma was dispersed throughout most of the frond area where generally it reached the edge, while in the other groups it was mostly in an upper-central position. Furthermore, only in a few individuals of the MG hybrid and *L. minor,* a centrally located aerenchyma was found. The papules trait also exclusively differentiated *L. gibba* from the other *Lemna* groups since papules were always absent in *L. gibba* and generally most evident in the MG cytotype, followed by *L. minor* and finally the GGM cytotype.

**Figure 8.**
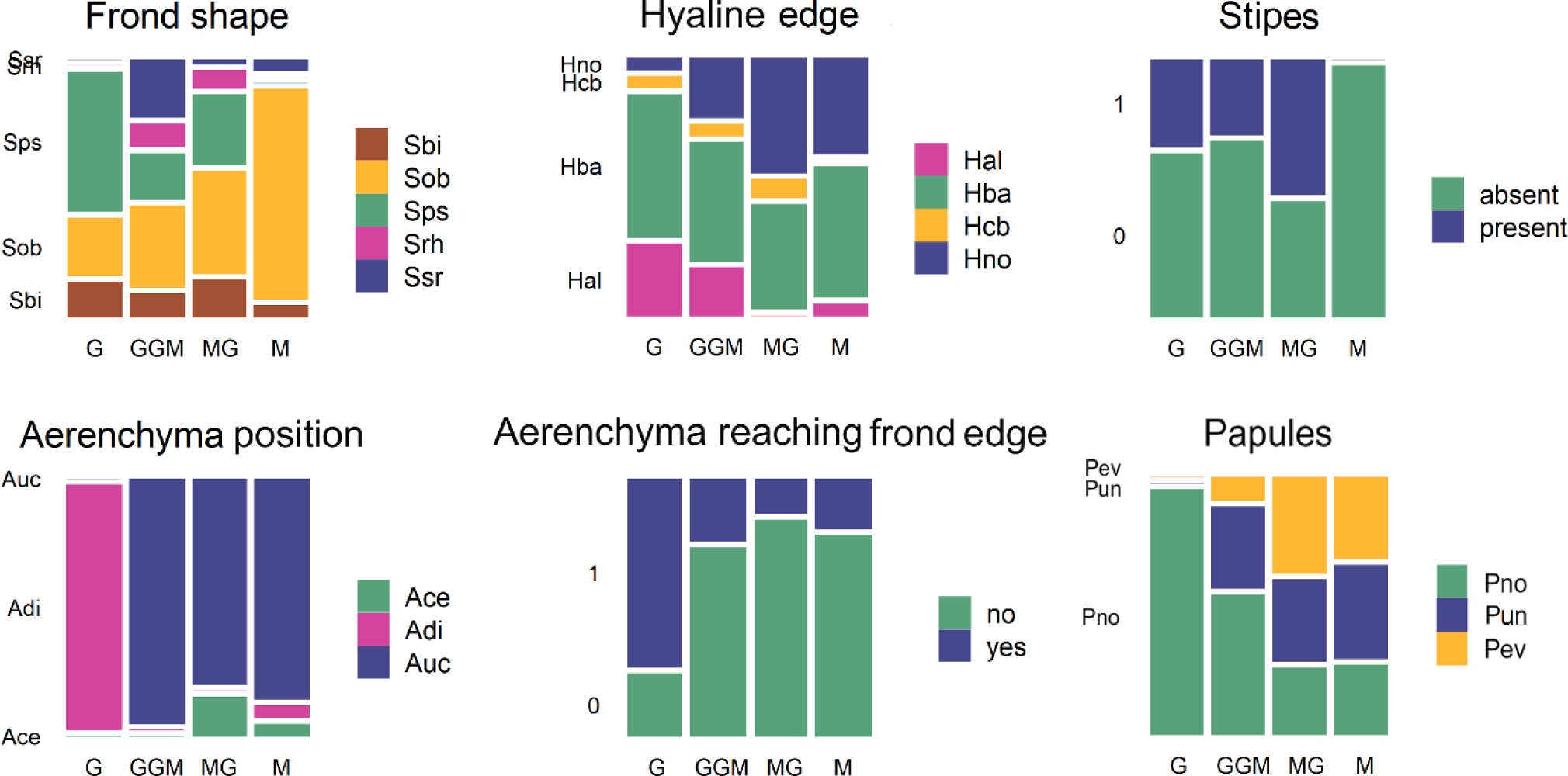
Differences among the four *Lemna* groups in relation to qualitative traits found to be significant by Chi-squared test (Mosaicplots). Acronyms for frond shape (Sbi = bilobate irregular shape, Sob = obovate, Sps = pear-shaped, Srh = rhomboid, Ssr = stocky rhomboid); hyaline frond edge (Hal = all around hyaline, Hba = basal, Hcb = central-basal, Hno = no hyaline); stipes present or not (absent); aerenchyma position (Ace = central, Adi = dispersed, Auc = upper-central aerenchyma); aerenchyma reaching (yes) or not (no) the frond edge; papules (Pno = absent, Pun = unclear, Pev = evident).

### Frond vascular organization

Differences in the simplified vascular tissues were observed comparing cleared frond specimens across the eleven *Lemna* clones considered. In Fig. 9, representative stereomicroscope images of fronds show visible interior veins within the body of the thallus. In particular, in *L. minor* 5500 a central vein and two lateral veins arising from the point of root attachment were present while in *L. gibba* 7742a five veins branched off from the node as reported in literature (Landolt, 1986; Bog, et al., 2019). MG hybrid cytotypes (e.g. 7641 and LM0027) exhibited from three to four veins per frond while GGM hybrid cytotypes predominantly revealed five veins as the maternal parent species

**Figure 9.**
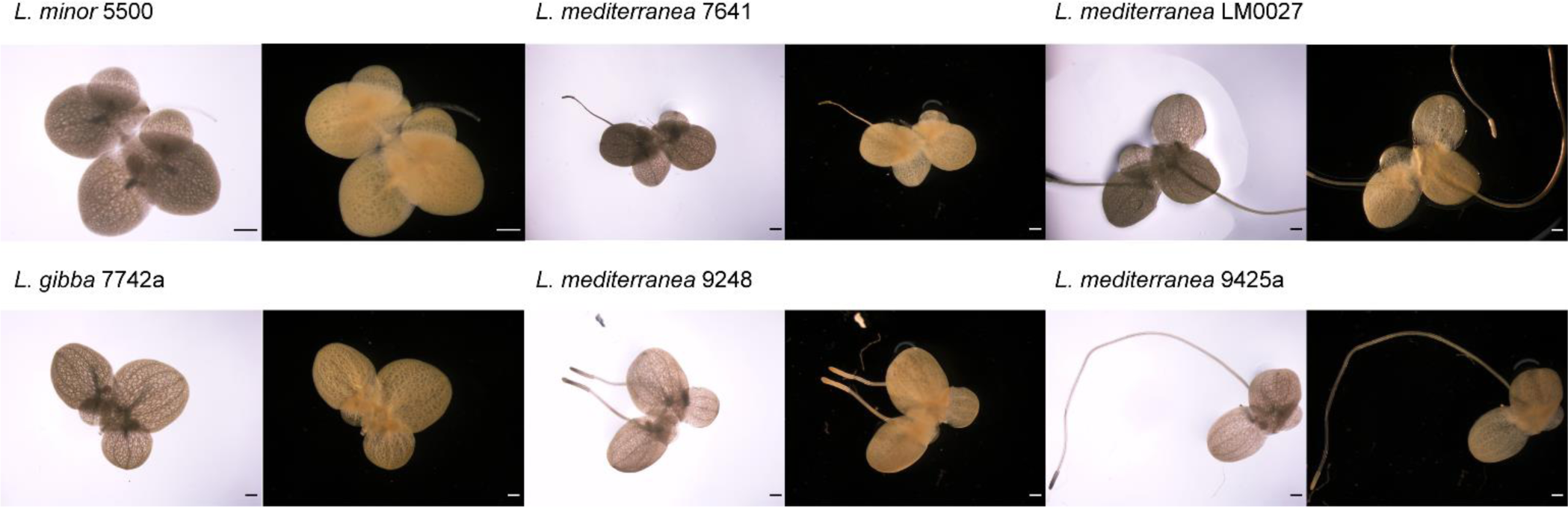
Representative stereomicroscope images of cleared fronds colonies for the determination of veins number per frond in the two parental species, *L. minor* 5500 and *L. gibba* 7742a, and in both *L. × mediterranea* cytotypes, homoploid (MG-7461 and -LM0027) and triploid (GGM-9248 and -9425a). Fronds colonies were observed under bright- and dark-field conditions. Bar = 1mm

### Stomatal traits

Stomatal density can be an indicator of the level of adaptation to environmental conditions. Stomatal size and density are dramatically impacted by growth environment factors, including light intensity, water stress, and CO2 concentration elevation. Measurement of stomatal size and density is summarized in Fig.10. In *L. minor*, stomatal density and size were correlated as the observed reduced stomatal density corresponded to a lower stomatal size. In particular, stomatal density and size in *L. minor* were significantly lower when compared to *L. × mediterranea* MG. The highest stomatal density was observed in the diploid MG. *L. gibba* and the triploid *L. × mediterranea* GGM did not significantly differ in stomatal density, also with respect to *L. minor* and MG. *Lemna minor* presented the smallest stomatal size, and GGM showed the highest. This is consistent with the fact that GGM clones have a higher DNA content than diploids, which usually correlates with cell size (McGoey et al., 2014; Da Silva et al., 2020).

**Figure 10.**
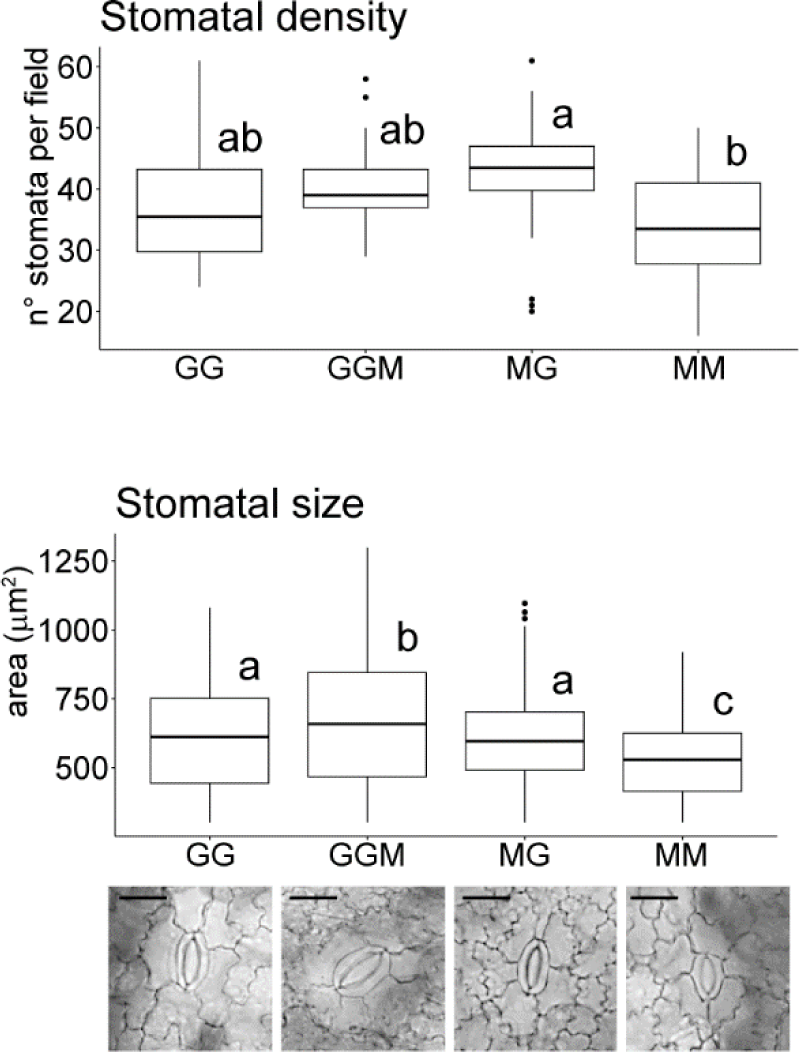
Differences in stomatal traits between the two parental species, *L. minor* (MM) and *L. gibba* (GG), and the two *L.* × *mediterranea* cytotypes, (MG and GGM): stomatal density (above) and size (below). Box plots labelled with different letters indicate significant differences between grouped *Lemna* species and hybrid cytotypes (ANOVA followed by Tukey HSD, p < 0.05). n = 3. Representative examples of stomatal morphology (bottom) in each of the corresponding groups photographed by optical microscopy. Bars = 30 µm

### Plant growth and biochemical characterization

The biochemical analysis (pigment and protein content), and plant growth parameters (RGR, frond fresh weight) showed that the two *L.* × *mediterranea* cytotypes exhibit their own independence and greater association to one of the parental species, as it is shown by PCA (Fig.11). PCA performed on the dataset captured 89.7% of the cumulative variance using the parameters influencing the first two principal components. The outcomes of PCA clearly discriminated *L. minor* (MM) and homoploid hybrid clones MG from *L. gibba* (GG) and triploid hybrids GGM. The profile of *L. minor* and MG clustered in a PC1-negative direction while *L. gibba* and GGM clustered in a PC1-positive direction.

**Figure 11.**
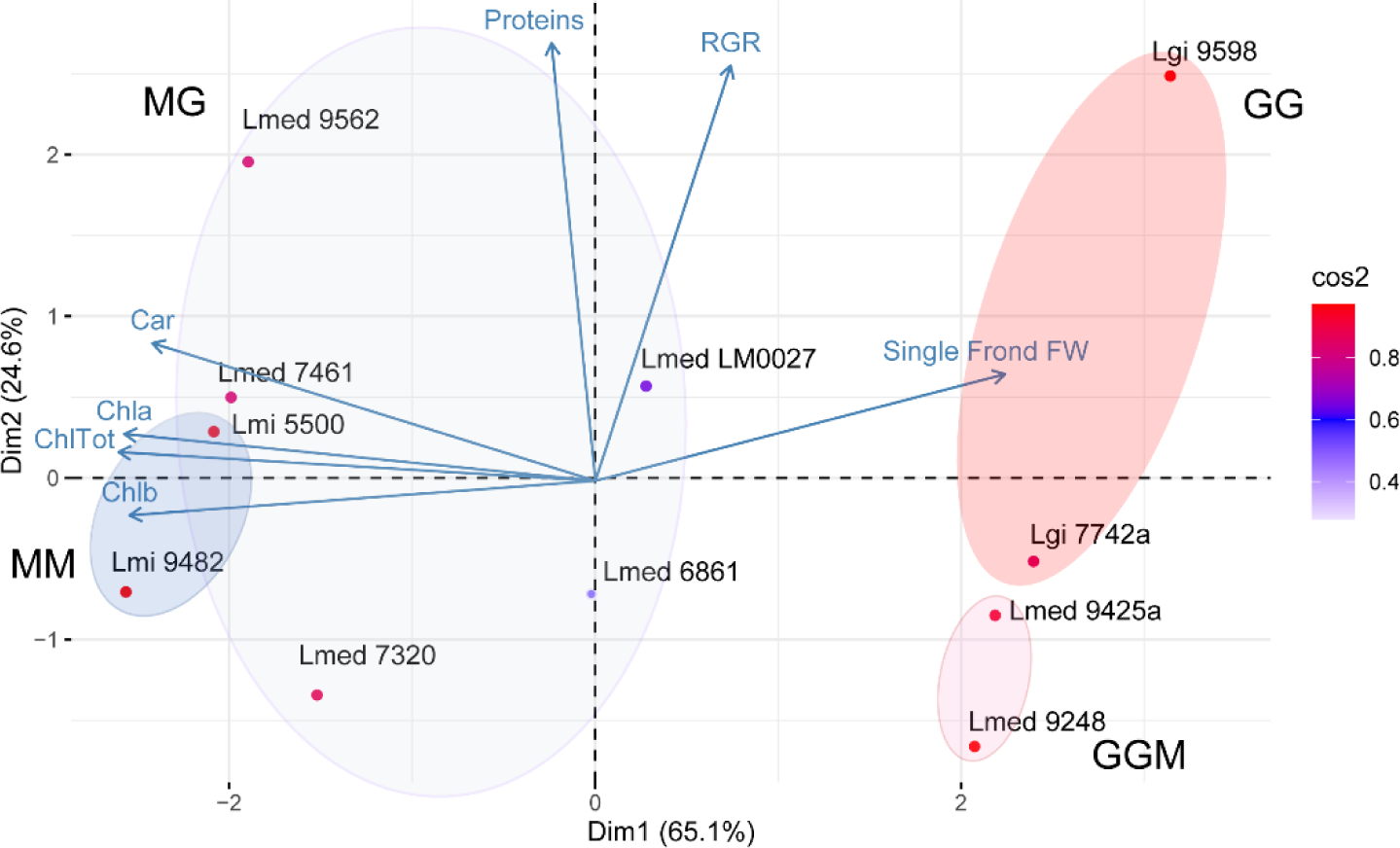
Principal component analysis (PCA) of the measured growth data and biochemical parameters: Relative Growth Rate (RGR), proteins, chlorophylls (ChlTot - Chla - Chlb), carotenoids (Car) and Single Frond Fresh Weight - (Single Frond FW) of the two parental species, *L. minor* (MM) and *L. gibba* (GG), and the two *L. × mediterranea* hybrid cytotypes (MG and GGM). Plot for PC1 and PC2, where each oval encompasses the observed pattern of variance of each *Lemna* clone under the first two principal components clustering separately MM and MG (blue ovals) and GG and GGM (pink ovals), respectively. Measured parameters are summarized as boxplots in Fig.12. No significant differences were found between *L. gibba* and triploid hybrids GGM in photosynthetic pigment content. The chlorophyll *a* content differed significantly between *L. gibba* and GGM hybrids compared to *L. minor* and MG hybrids, respectively. Furthermore, for chlorophyll *b* and carotenoid content significant differences were found between *L. minor* and hybrid cytotype homoploid MG and in respect to *L. gibba* and the hybrid cytotype triploid GGM. *Lemna minor* (*L. minor* 5500 and *L. minor* 9482) had the highest pigments content. The estimated fresh weight of single frond of *L. gibba* is significantly different with respect to *L. minor* and to both *L. × mediterranea* cytotypes. No significant differences were found between *L. minor* and the homoploid cytotype MG. Equally high growth rates under the tested conditions were shown inter- and intra- the two species, *L. minor* and *L. gibba* and the two hybrid cytotypes MG and GGM. In particular, RGR in the period under study ranged from 0.17 to 0.25 g^-1^ day^–1^for *L. gibba* and 0.17-0.18 g^-1^ day^–1^ for GGM clones, while 0.16 to 0.18 g^-1^ day^–1^for *L. minor* and 0.16 to 0.22 g^-1^ day^–1^ for MG. These RGRs values agree with data reported in literature, which are situated around 0.1 d^-1^ up to 0.3 d^-1^ (Zhang et al., 2014; Van Echelpoel et al., 2016). In the hybrid triploid cytotype, GGM, the protein content was lower and significantly different compared to the homoploid cytotype MG and to the two parental species. Among the analysed accessions, *L. gibba* 9598 showed the highest values for single frond fresh weight, RGR and protein content.

### Taxonomy

A natural interspecific hybrid between *L. minor* and *L. gibba* is described here:

***Lemna* × *mediterranea*** L. Braglia & L. Morello, hybrida nova, *L. minor* × *L. gibba*

**Type**: Italy. Umbria Region, Passignano on the Trasimeno Lake (4310.948, 1209.297, 257 m elevation) [clone 9562], collected in 2011 by *F. Landucci*. Fig. 13a, b.

**Figure 12.**
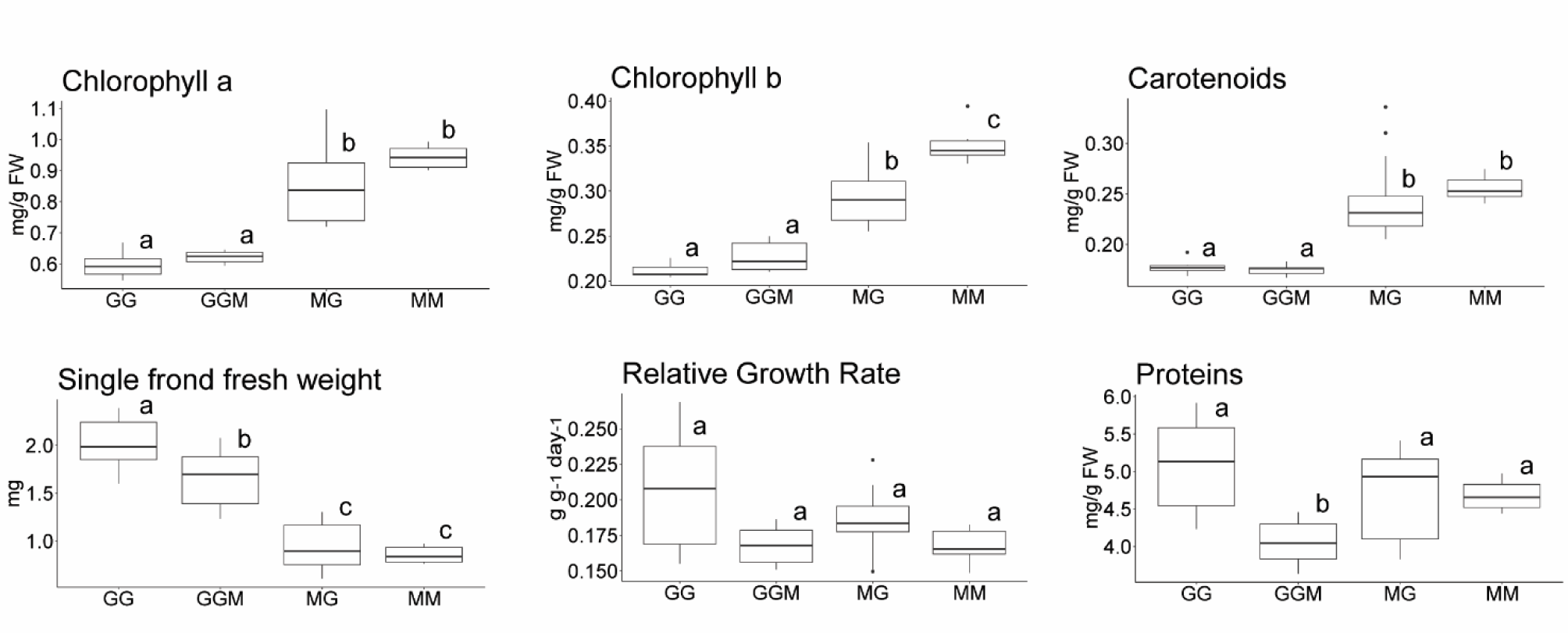
Differences in photosynthetic pigment content, single frond fresh weight, relative growth rate (RGR) and protein content between the two parental species, *Lemna minor* (MM) and *L. gibba* (GG), and the two *L. × mediterranea* hybrid cytotypes MG and GGM. Box plots labelled with different letters indicate significant differences between different grouped *Lemna* species and hybrid cytotypes (ANOVA followed by Tukey HSD, p value ≤ 0.05). n = 5

**Figure 13.**
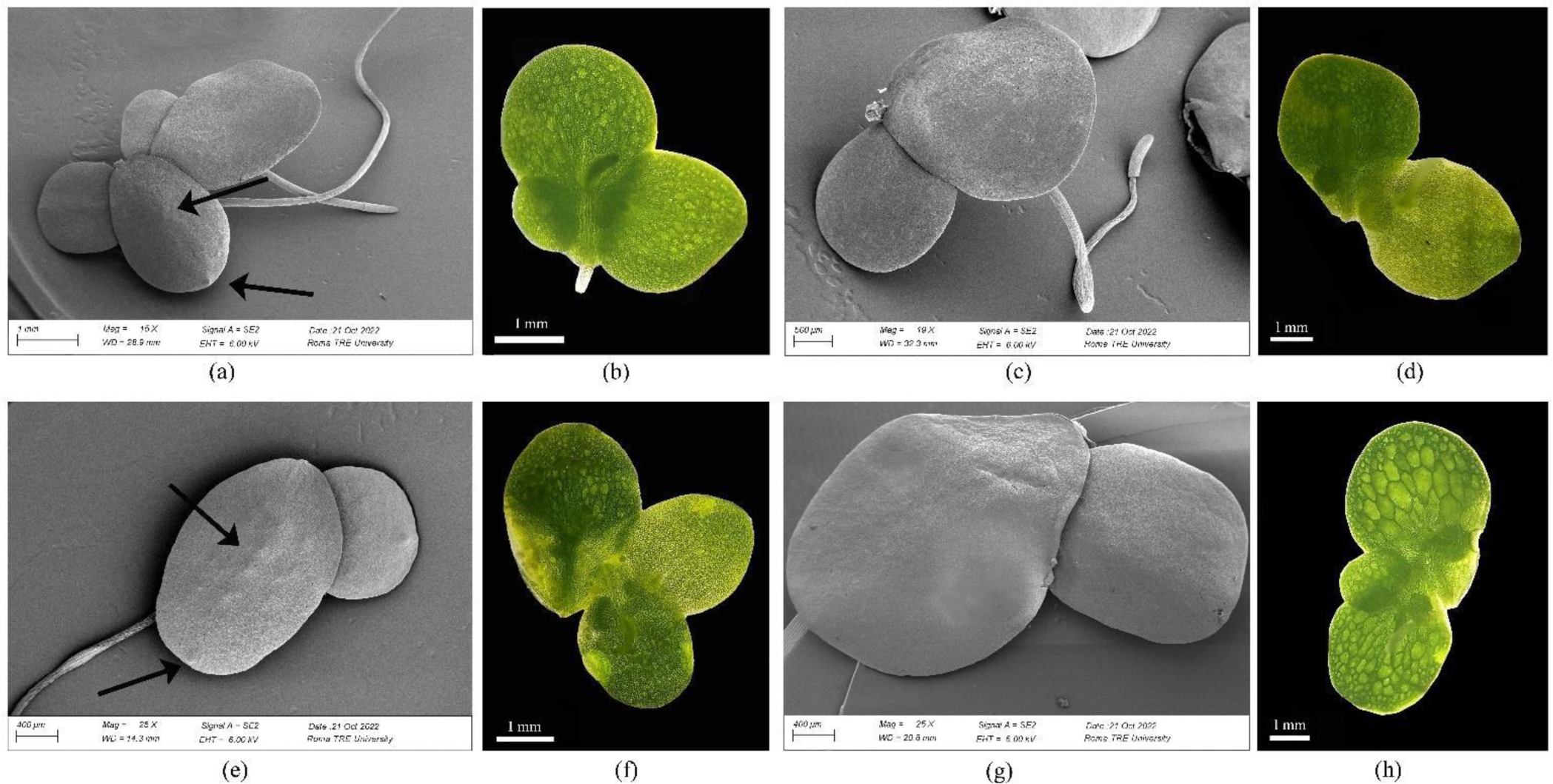
Representative images of the hybrids *L. × mediterranea* - homoploid cytotype (MG), clone 9562 (a-b) *L. × mediterranea* - triploid cytotype (GGM), clone 9425a (c-d) and the parental species, *L. minor* (e-f), *L. gibba* (g-h) at scanning electron microscopy (left) and stereoscope (right). In detail: adaxial frond surface (a, c, e, g) and abaxial frond surface with visible aerenchyma (b, d, f, h). Dark arrows indicate serial or terminal papules on the adaxial frond surface (a, e).

Specimens of the holotype as well as all the other six clones described in the present paper are being deposited at the Central Herbarium of Italy, at the University of Florence. Specimens of the same clones, except for LM0027, are also present in a large herbarium on duckweeds assembled by Prof. E. Landolt († 1921-2013) which is presently held privately in Zurich by Mr. W. Lämmler.

**Synonym:** ‘*Lemna symmeter’* G. Giuga [Giuga G., 1973; “Vita segreta di Lemnacee I. *Lemna symmeter* G. Giuga-species nova”, Tip. Di Blasio, Naples], pro sp., nom. non rite publ. (nec descr. Lat., non typ.). Supplementary Figure 4. This name was previously considered as a synonym of *L. gibba* L. (Sree et al., 2016). Moreover, as no Latin description was present, it was not effectively published according to the requirements of the *International Code*. In any case, ‘*L. symmeter’* is not grammatically correct Latin (the correct adjective should have been *symmetrica*, meaning symmetric). No authentic specimens of ‘*L. symmeter’* have been recorded in any official herbaria to our knowledge, so this synonymy cannot be unquestionably ascertained.

### Morphological description

Morphological characters are intermediate between the two parental species *L. minor* and *L. gibba*, with greater morphometric similarity to *L. minor.* As for frond shape, this hybrid shows a variable morphology, even if the obovate shape is slightly more frequent than others. Fronds have an average length of 3.60 mm (2.71– 4.87 mm), width of 2.54 mm (2.04 – 3.07 mm) and an area of 6.53 mm^2^. No evident gibbosity is observed. The number of visible veins is from 3 to 4. Similar to *L. minor*, several serially arranged papules are often visible along the mid vein on the adaxial frond surface. Sometimes there is an additional, prominent, isolated papule positioned near the tip of the frond. The aerenchyma is mainly localized in the mid-upper part of the frond, covering on average ca. 54.60% of the frond area. Average length of the longest aerenchyma cells is 0.23 mm (0.13–0.38). The mean root length is 2.81 mm with a maximum value of 6.91 mm. Over 50% of the analysed fronds showed elongated stipes connecting daughter fronds.

**Ploidy:** as inferred by comparison of the relative genome size with that of the parental species, the hybrid type is homoploid (MG). Karyotyping is needed for full confirmation.

**Lemna × mediterranea** – reversed cross *L. gibba* × *L. minor* (Fig. 13 c-d)

Morphological analysis of the two clones genetically attributable to this cytotype shows that it also exhibits intermediate characters between the two parental species, while showing greater morphological similarity to *L. gibba*, in accordance with the double genetic contribution of this latter. In this hybrid, the frond shape is variable, not showing one shape predominant over the other; fronds are larger than the diploid, with an average length of 3.74 mm (3.24 - 4.82), width of 2.87 mm 2.56 - 3.26), and total area of 7.90 mm^2^. No evident gibbosity is observed. The number of visible veins is predominantly 5. On the adaxial frond surface, some serially arranged papules are sometimes visible in a hinting manner. Aerenchyma tissue is developed mainly in the mid-upper portion of the frond and covers on average 72.50% of the area, never reaching the frond margin. Average aerenchymal cell length is 0.25 mm (0.15 - 0.43). The mean length of roots is 5.87 mm with maximum value of 11.15 mm. In 30% of the samples analysed, the presence of elongated stipes connecting neighbouring fronds was observable at the base of the frond. Guard cells are larger than in the diploid cytotype.

**Ploidy:** both clones representing this cytotype are triploid, with two subgenomes acquired from the maternal parent *L. gibba* (GGM), as deduced from qPCR analysis. Karyotyping will provide further confirmation.

**Diagnosis (Recognition):** Because of the wide phenotypic plasticity, the existence of two different cytotypes, and the presence of intermediate morphological traits with respect to the parental species, no dichotomous key can be developed for the straightforward recognition of the hybrid *L.* × *mediterranea*. Although all hybrid clones in the original Landolt collection were classified as *L. gibba* by morphology, the morphometric analysis performed here showed a closer overall similarity of the homoploid hybrid to *L. minor*, and the triploid hybrid to *L. gibba*. As morphological recognition of *L.* × *mediterranea* is almost impossible, diagnosis must then rely on molecular markers. Since plastid markers are of no help, we suggest tubulin intron amplification (*TUBB*1 and *TUBB*2) followed by agarose gel electrophoresis as a very simple molecular method for fast identification, as described in Braglia et al., 2021b. Such analysis is suggested every time a *Lemna* specimen cannot be assigned with accuracy to either *L. minor* or *L. gibba*.

**Distribution:** The geographical origin of the seven hybrid specimens is reported in Fig. 14. All but one of them, which was sampled in the North of Germany, come from Mediterranean countries: four from Italy (at different latitudes), one from Israel and one from Egypt.

**Figure 14.**
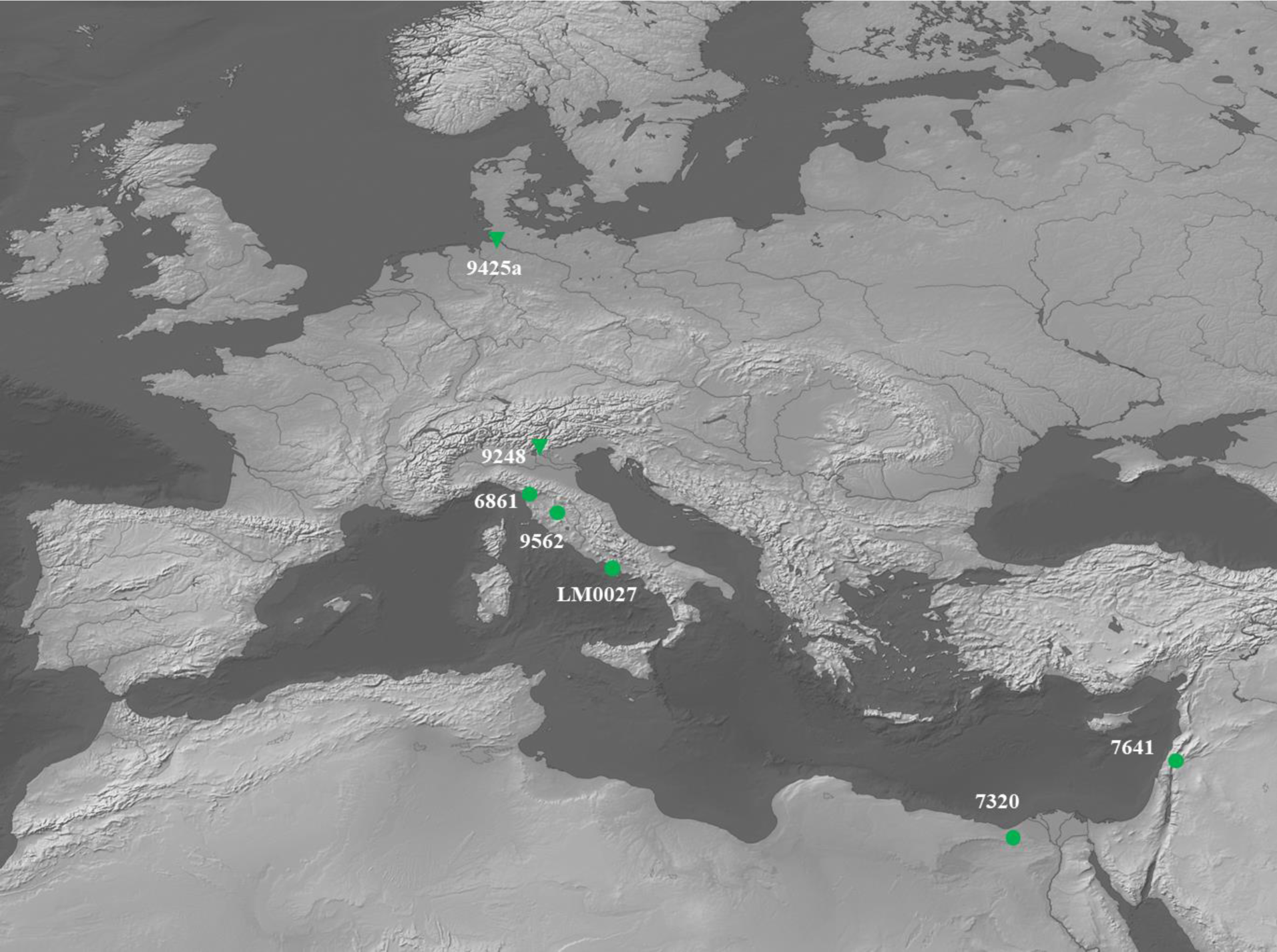
Geographical origin of the seven *L. × mediterranea* specimens. Circles indicate homoploid hybrids MG, triangles indicate triploid GGM clones.

Among a total of 23 *L. gibba* and 48 *L. minor* specimens from our present collection and coming from all continents, no other hybrid clone was found. This fact suggests a rather restricted distribution of *L.* × *mediterranea* despite wide overlaps in the geographic distribution of *L. gibba* and *L. minor* and the co-occurrence of the two species in the phytosociological alliance *Lemnion minoris* at many sites of Europe, South-western Asia, Africa and California (Landolt, 1986).

**Etymology:** The specific epithet refers to the collection sites of six out of the seven investigated specimens, coming from three different Mediterranean countries

**Conservation status**: Not known. All clones investigated were from *ex situ* germplasm collections. However, collection dates of the seven clones span nearly 70 years, from 1954 to 2006, suggesting either recurrent hybridization or population stability. Field research is ongoing in some of the areas where hybrid clones were collected.

**Fertility:** Not known. Flowering of ‘*L. symmeter’* was reported as common by G. Giuga, both in the wild and in quasi-natural conditions (samples collected in the wild and grown in water outdoors), producing indehiscent, sterile anthers (Giuga, 1973). This agrees with the fact that both genomic arrangements, homoploidy and triploidy, are generally associated with the absence or a severely reduced fertility in other plant species (Rieseberg et al., 2007). Trials to induce flowering are ongoing to confirm sterility.

### Morphological description of the parental species

For comparison, we provide a description of the parental species. Not all the features reported are in full agreement with those described in literature as our description is limited to few European clones, since long cultivated *in vitro*.

***Lemna minor*** (Fig. 13 e-f)

Fronds of *L. minor* show a predominantly obovate shape (never pyriform), averagely 3.4 mm (2.5 – 3.9 mm) long and 2.4 mm (1.75 – 3.0 mm) wide, with an area of 6.0 mm^2^ (3.7–8.45 mm), on average smaller than *L. gibba*. The number of veins is predominantly 3. Several serially arranged papules are often visible, sometimes prominently, on the adaxial frond surface. Often there is an additional isolated papule near the frond tip (Fig. 13e). Aerenchyma is localized in the mid-lower part of the frond and is covering on average 40% of the frond area. Average length of aerenchyma cells is 0.16 mm (0.10–0.33 mm). The maximum root length recorded was 11 mm. Absence of stipes connecting contiguous fronds within a cluster.

***Lemna gibba*** (Fig. 13 g-h)

Based on morphological analysis, *L. gibba* is characterized by having predominantly pyriform fronds, on average 3.9 mm long (2.6–5.1) and 3 mm wide (2.3–3.8), with a total area of 8.5 mm^2,^ larger than *L. minor*. The number of visible veins is 5. No papules are visible on the adaxial frond surface. Highly developed aerenchyma tissue covering on average more than 90% of the frond area and generally reaching its edge. Average length of aerenchyma cells, 0.53 mm (0.26–0.88). Maximum measured root length was 35.4 mm. Sometimes (35%), formation of elongated stipes connecting daughter to mother fronds.

## DISCUSSION

Multiple molecular data provided definite evidence that *L. minor* and *L. gibba* can spontaneously hybridize in nature, confirming previous data obtained by TBP analysis (Braglia et al., 2021). An example of hybridization between these two species is here given by the first description of the new nototaxon, *Lemna* × *mediterranea*, up to now overlooked. This hybrid likely corresponds to the taxon described (not validly) as the new species ‘*L. symmeter’* by G. Giuga (G. Giuga, 1973), although its hybrid nature was not suspected by the author. Hybridization was instead hypothesized by Kandeler (Kandeler, 1975) while reporting that the proposed taxon *L. symmeter* stands between *L. gibba* and *L. minor* and might even be a sterile hybrid between these two *Lemna* species. The description of *L.* × *mediterranea* underscores how in this hybrid there is the appearance of intermediate morphological traits between the two parental species, a common event in hybrids that often makes their morphological discrimination from the parental species challenging. Despite limited morphological differences between hybrids and parental species, morphometric analysis of several traits is in agreement with genetic analysis in supporting a clear distinction of *L* × *mediterranea* and also in separating the two cytotypes, homoploid (MG) and triploid (GGM). Paradoxically, each of the two cytotypes is more similar to one of the parental species than to the reciprocal hybrid. Whether this could be actually considered as a maternal effect or a gene dosage effect remains to be established.

The first cytotype is more closely related to *L. minor* while the second to *L. gibba*. The more distinctive morphological differences between the two cytotypes are mainly related to quantitative traits (frond width, frond area, length/width ratio, root length, aerenchyma extension, vein number) and only secondarily to qualitative ones (hyaline frond edge, aerenchyma position). Analysis of stomatal morphological parameters (size and density) highlighted the presence of significant differences in guard cell size, which are the largest in the GGM cytotype. This is likely related to its increased genome size, as already reported for *Lemnaceae* (Hoang et al., 2019). Also for the biochemical traits observed, as pigment content and RGR, hybrids have intermediate values, with triploid hybrids more closely related to *L. gibba* and diploid clones more similar to *L. minor*, suggesting that different genome contributions also affect biochemical traits and, possibly, plant physiological performance. Higher ploidy may also enable enhanced genetic capacity to respond to abiotic stress that is worth to be investigated. In conclusion, no morphological criterion is *per se* sufficient to provide unequivocal identification of *Lemna × mediterranea* clones, and the use of molecular analysis is strongly suggested.

As from the analysed samples, *L.* × *mediterranea* is distributed over a wide geographic area, centred in the Mediterranean region, and includes both reciprocal crosses of *L. minor* and *L. gibba*, as revealed by plastid marker analysis. Population structure analysis inferred from AFLP data using different bioinformatics models, subtend the occurrence of different lineages, the parental populations, converging in the formation of an interspecific hybrid population. In this respect, the limited number of private alleles detected in *L.* × *mediterranea* suggests a fully and bipartisan genomic contribution of both parents merged in the hybrid. Phylogenetic network reconstruction also identifies the dual contribution of the parent species, placing the hybrid group closer to *L. gibba* than to *L. minor*, and supports its separation in at least two, possibly three, diverging clusters (Fig. 4), also in agreement with the similarity tree generated from TBP profiles (Fig. S2). Greater similarity of hybrids to European clones of the parent species suggests their origin from a limited number of European ecotypes, in accordance with their lower intrataxon genetic diversity with respect to parental species. Some degree of genetic diversity among hybrid clones, although limited, favours the interpretation that each clone originated independently from different parental clones. In fact, according to NewHybrids results, all clones have high probability to represent F1 populations. No evidence for backcrossing emerged for the seven clones, despite *L.* × *mediterranea* may occur in association with either of the parental species, as reported in Italy for *L. symmeter* (Giuga, 1973) and for clones identified as non-gibbous forms of *L. gibba* but having the plastid haplotypes of *L. minor* (Marconi et al., 2019). Such observations suggest sterility or very low fertility and self-incompatibility of hybrids. Flower induction experiments are ongoing to address these key points. However, given the low frequency of flowering of the parental species, particularly *L. minor* (Landolt, 1986), we cannot expect inbreeding or outcrossing to be frequent events in hybrids so that the possibility of producing hybrid, self-evolving lineages slowly leading to speciation, cannot be excluded. Even very low rates of sexual reproduction are in fact considered sufficient to get rid of negative mutations that accumulate in asexual populations (Hojsgaard and Hörandl, 2015). In such a framework the observed diversity can be interpreted as the result of somatic mutation accumulation in long lasting asexual lineages. Although aggregates of vegetative reproducing individuals are unlikely to establish species-like lineages (Hörandl, 2022), in the case of homoploid hybrids, speciation is now accepted even if the hybrid lineage can be established as viable progenies through vegetative (or clonal) propagation, not necessarily requiring allopolyploidisation (Comai, 2005; Sochor et al., 2015; White et al., 2018). In *Lemna*, hybrid population stability and diffusion can be clearly provided by fast clonal propagation and long-distance dispersal of these tiny plants through water flow and zoochory (Coughlan et al., 2017), promoting its establishment as a species if favoured by some competitive advantage with respect to the parental species. The success of *L.* × *mediterranea* is evidenced by the large geographic area and collection dates of the hybrid clones from 1954 to 2011. Recovering living populations will provide further information on hybrid distribution and origin.

Another peculiarity of *L.* × *mediterranea* is the presence of two different but unusual cytotypes, homoploids with *L. gibba* as the pollen donor and triploids with *L. gibba* as the mother parent, while no tetraploid was found among hybrids until now. As both parental species are known to be mostly diploids (Landolt, 1986), the simplest explanation is that triploid hybrids originated from the fertilization of unreduced *L. gibba* ovules (2n) by normal haploid pollen cells (n) from *L. minor*. However, breeding between a tetraploid *L. gibba* and a diploid *L. minor* cannot be excluded. A somatic mutation leading to tetraploidy has been recently described for a *L. gibba* clone since long cultivated *in vitro* (Sarin et al., 2023). Wide variations in both genome size and chromosome number have been often reported in *Lemna* and *Wolffia*, although not all old chromosome counting data are fully reliable (Hoang et al, 2018; Hoang et al., 2022). More recent data show that triploid cytotypes are present in both *L. minor* and in the hybrid species *L.* × *japonica* that also includes homoploid hybrids (Ernst et al., 2023; Abramson et al., manuscript in preparation). The situation of the two intraspecific *L. minor* hybrids is in fact very similar, although *L. japonica*, recognized as a species in 1980 (Landolt, 1980), has a larger geographic distribution in the northern hemisphere, from Eurasia to North America, and a wider intraspecific variability (Braglia et al., 2021b) in comparison with *L.* × *mediterranea*. Also in that case, self-fertility and seed production has not been ascertained. In both cases, the question if these large hybrid populations should be considered as true or potential species requires further investigation.

Hybridization is extremely common in plants and most successful hybrids are polyploid, a condition which grants full fertility eventually leading to hybrid speciation. Conversely, both homoploid and triploid hybrids are quite rare in terrestrial plants and are considered as bridges to form fully fertile, higher ploidy (tetraploid/hexaploid) species (Ramsey and Schemske 2002). In a very few cases, homoploid hybrids become stabilized over time, keeping an acceptable degree of fertility and becoming reproductively isolated from parents thanks to ecological or biological barriers (Mason and Pires, 2015) then becoming morpho-physiologically different, self-evolving species. Homoploid hybrid speciation (HHS) has been well documented in some plant species as *Helianthu*s (Schwarzbach 2002), or *Senecio* (Abbott et al., 2013), but true numbers are likely underestimated (Yakimoski and Rieseberg, 2014). The number of known triploid plant species is even smaller, partially due to the triploid block effect, impairing endosperm development and inducing seed abortion (Kohler et al., 2010). In such cases, at least at early stages, clonal propagation can provide an escape route to the low degree of fertility (Vallejo-Marin and Hiscok, 2016). An interesting example of a recently generated triploid species, *Cardamine × insueta* Urbanska-Worytkiewicz, have been documented in the Swiss Alps (Urbanska and Landolt, 1972). The colonization of a new habitat provided almost completely reproductive isolation from the parental species while the acquisition of leaf vivipary enabled the hybrid to be a dominant species at the site despite its ploidy level (Sun et al., 2020). More detailed analysis of ecological differences between *Lemna* hybrids and parental species is also needed to understand the advantages of hybrids and the possibility of their adaptation to different ecological niches even within the same water body.

This study further demonstrates that interspecific hybridization can be a common mechanism to generate diversity and variation in *Lemna*, which might have played an important role in the evolution and diversification of this genus and, possibly, in other genera of duckweeds. This is in accordance with findings by Les and Philbric (1993) who, analyzing literature data for 117 genera of aquatic angiosperms, suggested that the high vagility (displacing ability) and rarity of sexual reproduction common to most of them has dramatically influenced the evolutionary consequences of two factors that have played major roles in the evolution of terrestrial angiosperms, namely hybridization and chromosome number change.

## Author Contributions

Conceptualization, L.M., S.C, M.A.I.; duckweed clone maintenance and investigation, M.A.I, G.F., M.F, S.C., F.M., E.P., L.B.; formal analysis, M.B., E.P., M.A.I, G.F, L.B.; data curation M.M.; writing original draft preparation, S.C, M.A.I, L.M, M.B.; writing review and editing, L.B, M.B., S.C., M.A.I.; G.F., S.G.; visualization, M.B, M.A.I, G.F.,M.F, F.G., S.C, E.P., M.M., L.B; supervision, L.M.; funding acquisition, S.C, M.A.I, L.B, L.M.

All authors have read and agreed to the published version of the manuscript.

## Data Availability Statement

Not applicable.

## Acknowledgments

We wish to warmly thank Diego Breviario for the fruitful TBP legacy and Walter Lammler for providing clones of the Landolt’s collection

