## Supplemental files for "Characterization of the cryptic interspecific hybrid *Lemna × mediterranea* by an integrated approach provides new insights into duckweed diversity"

**SUPPLEMENTARY DATA**

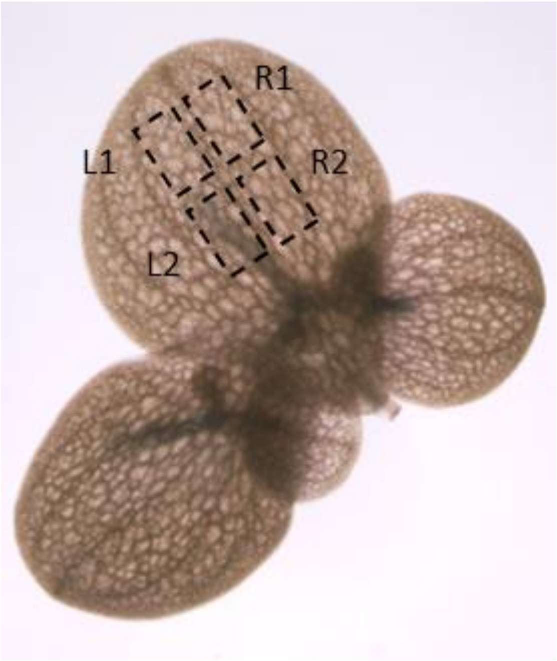


**Figure S1.** Image example used for measuring stomatal traits in the four different microscopic fields identified by two sectors on each side (L1, L2 and R1, R2) of the mother frond main vein.

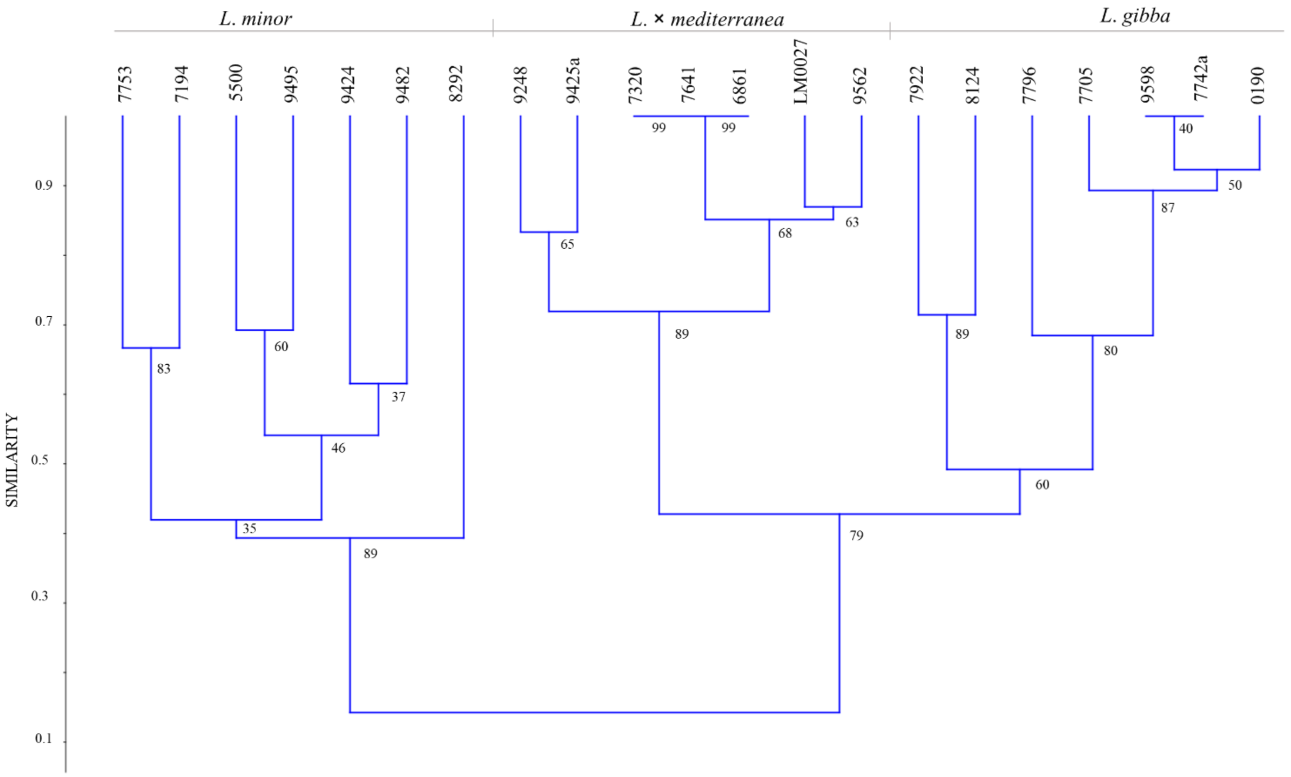


**Figure S2**. Dendrogram generated by UPGMA clustering analysis of the TBP dataset of all analysed *Lemna* clones. Numbers at nodes refer to bootstrap values (in %) estimated for 1000 replicates.

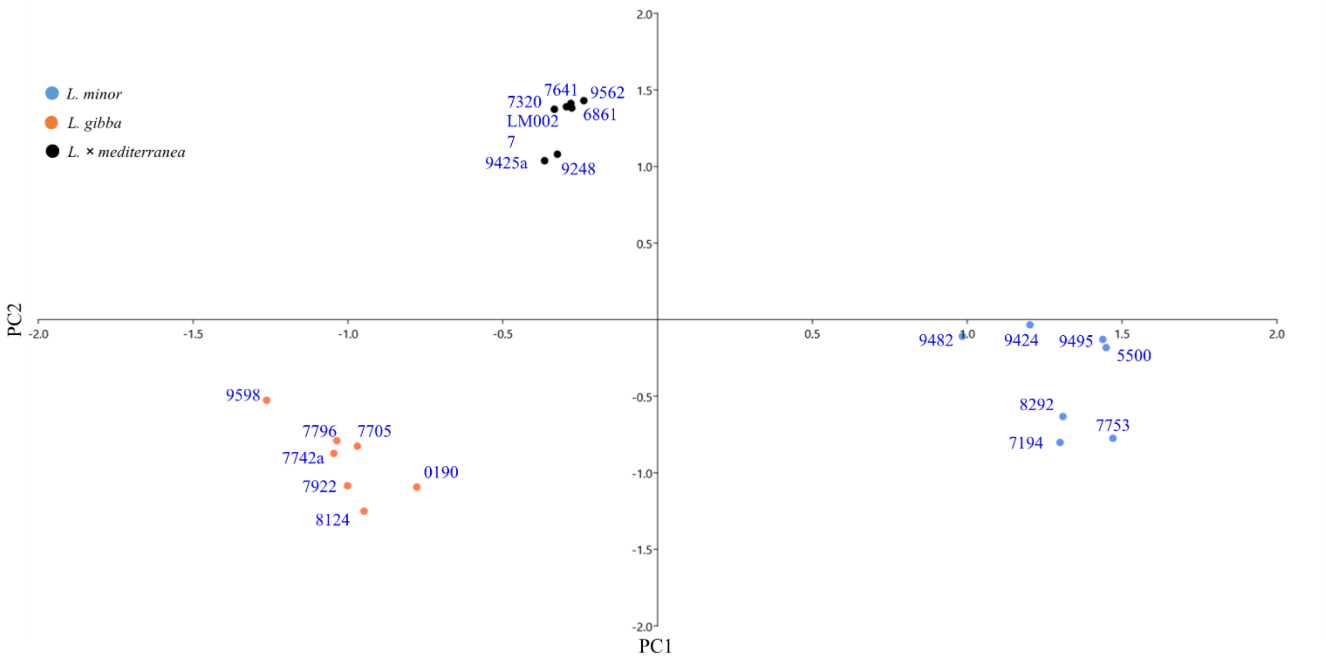


**Figure S3.** PCA of the AFLP dataset of *Lemna* clones analysed. Total variance (PC1 37% - PC2 17%)

G. GIUGA

### Vita segreta di *Lemnacee*.

#### *Lemna symmeter* G. Giuga - species nova.

Tip. Di Blasio - Via Costantinopoli, 104 - 80138 Napoli - Tel. 452508
1973



SECRET LIFE OF *Lemnaceae*. *Lemna symmeter* G. Giuga species nova

SUMMARY

The new species, *L. symmeter*, at present is widely distributed on Campania coast-line, in the several sunny canals of the widening Low Volturno large alluvial basin, or in those of Sarno smaller basin.

It is associated with *L. gibba*, much more common, characterized by a population of globose, almost spheroidal fronds, which flowers every, with fruitful seeds, in the most sunny canals, *L. symmeter* is instead, lightly *ventricous* and generally sterile. In *L. gibba* the pollination process isn't clear (W. S. Hillman, 1961).

It was necessary to clarify and complete it by observing over and over again a *primitive* cultivation of *L. gibba* led to flowering. The results obtained by causing the coming out of the wole inflorescences using sub-microscopic methods and by observing them at particular stages, seemed to us to be like the results obtained, as regards the flowering of *L. paucicostata*, by the eminent embryologists S. C. MAHESHWARI and R. N. KAPIL J1963) *L. gibba* and*L. paucicostata* (placed in synonymy with the American *L. perpusilla*), are the only species of *Lemna* that more often flower with fertile seeds.

*L. gibba* flowering process can be described as follows:

Since the first stages, the two anthers develop typically out phase, asymmetrically, and arise from the pouch *minus* (opposed to that in which the first daughter frond is produced) at different times.

Usually, the pistil ripes first; anymay the pistil and the first anther appear almost simultaneously, very near each other. The second anther when it is near the flowering, is sometimes smaller than the first one.

In the new species *L. symmeter* the inflorescence develops in a very different way:

First the stigma arises solitary from the pouch *minus*. At this stage, and during several days when the stigma remains solitary, the two onthers enclosed in the pouch where it was produced, grow symmetrically, sub-equal.

Later, the two anthers, arise undehisced from the pouch minus, whereas the stigma no longer receptive withers.

All the floral parts become disorganized.

The new species is generally sterile.

At the vegetative stage, *L. symmeter* three morphological characters have been sufficient for individualizing the species:

1) almost sym metrical fronds whith a rounded apex and
2) lightly *ventricous;*
3) regular air spaces smaller than *L. gibba* ones.

W. S. HILLMAN, (1961); The *Lemnaceae* Bot. Revue **27**, 271-410.

S. C. MAHESHWARI e R. N. KAPIL, (1963; Morphologycal and Embriological studies of lemnacee I *The floral structure of the gamotophytes of Lemna paucicostata*. (Am. Jour. Bot. **50** (7); 677-688).

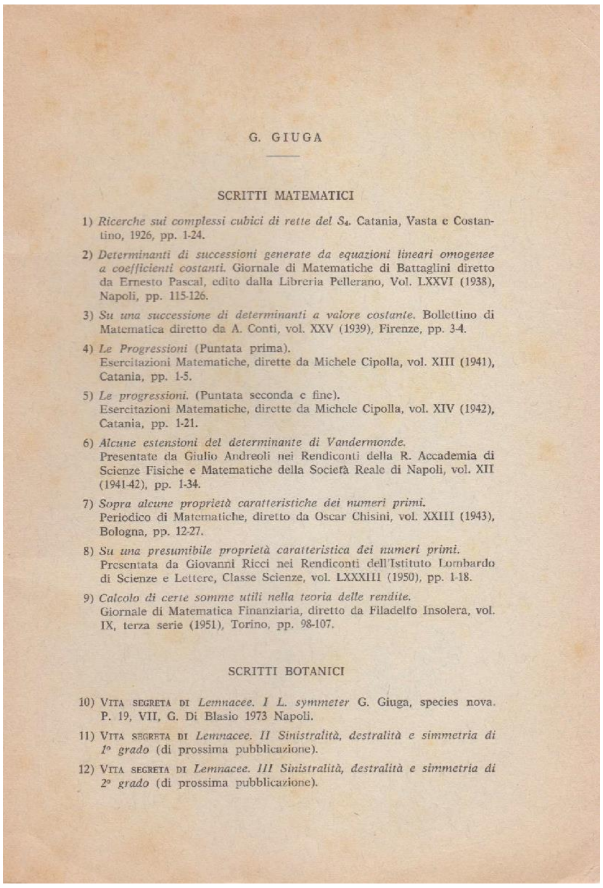

G. GIUGA

SCRITTI MATEMATICI

1) *Ricerche sui complessi cubici di rette del $S_4$.* Catania, Vasta e Costantino, 1926, pp. 1-24.

2) *Determinanti di successioni generate da equazioni lineari omogenee a coefficienti costanti.* Giornale di Matematiche di Battaglini diretto da Ernesto Pascal, edito dalla Libreria Pellerano, Vol. LXXVI (1938), Napoli, pp. 115-126.

3) *Su una successione di determinanti a valore costante.* Bollettino di Matematica diretto da A. Conti, vol. XXV (1939), Firenze, pp. 3-4.

4) *Le Progressioni* (Puntata prima).
Esercitazioni Matematiche, dirette da Michele Cipolla, vol. XIII (1941), Catania, pp. 1-5.

5) *Le progressioni.* (Puntata seconda e fine).
Esercitazioni Matematiche, dirette da Michele Cipolla, vol. XIV (1942), Catania, pp. 1-21.

6) *Alcune estensioni del determinante di Vandermonde.*
Presentate da Giulio Andreoli nei Rendiconti della R. Accademia di Scienze Fisiche e Matematiche della Società Reale di Napoli, vol. XII (1941-42), pp. 1-34.

7) *Sopra alcune proprietà caratteristiche dei numeri primi.*
Periodico di Matematiche, diretto da Oscar Chisini, vol. XXIII (1943), Bologna, pp. 12-27.

8) *Su una presumibile proprietà caratteristica dei numeri primi.*
Presentata da Giovanni Ricci nei Rendiconti dell'Istituto Lombardo di Scienze e Lettere, Classe Scienze, vol. LXXXIII (1950), pp. 1-18.

9) *Calcolo di certe somme utili nella teoria delle rendite.*
Giornale di Matematica Finanziaria, diretto da Filadelfo Insolera, vol. IX, terza serie (1951), Torino, pp. 98-107.

SCRITTI BOTANICI

10) VITA SEGRETA DI *Lemnacee. I L. symmeter* G. Giuga, species nova. P. 19, VII, G. Di Blasio 1973 Napoli.

11) VITA SEGRETA DI *Lemnacee. II Sinistralità, destralità e simmetria di 1° grado* (di prossima pubblicazione).

12) VITA SEGRETA DI *Lemnacee. III Sinistralità, destralità e simmetria di 2° grado* (di prossima pubblicazione).

**Figure S4.** Giuga's Booklet: The secret life of Lemnaceae. His original publication, in Italian, is cited in the literature by Kandeler (1975) and Landolt (1986) but we could not find it in any public library or online repository. Curiously, we were able to recover one copy of the original monograph from a used bookseller in Naples (Italy), by buying it online. This allowed us to learn that G. Giuga was not a botanist, rather a mathematician, as witnessed by some of his mathematical works mentioned in the booklet. We realised that he is remembered for a theory on prime numbers that is known as Giuga's conjecture (Giuga, 1950). A second and third issue of the monograph, dealing with symmetry in *Lemna*, were planned but perhaps were never printed.

**Supplementary Table 1.** Oligonuclotide list

| | Oligonucleotide | Sequence(s) |
|---|---|---|
| Homoeolog-specific qPCR | TUBB2 minor_specF | GAAAACTGCGACTGCCTCCA |
| | TUBB2 gibba_specF | GGAAAACTGCGACTGCCTTCA |
| | TUBB2 gibba_specR | CAAGGATGAGAATAGATCGCAAC |
| | TUBB2 gibba_specR | TCGGCAAGGATGAGAATATTGC |
| AFLP Adaptors | *Mse*-Ad 3’ | GACGATGAGTCCTGAG |
| | *Mse*-Ad 5’ | TACTCAGGACTCAT |
| | *Eco*-Ad 5’ | AATTGGTACGCAGTCTAC |
| | *Eco*-Ad 3’ | CTCGTAGACTGCGTACC |
| AFLP Pre-amplification | *Mse02* | GATGAGTCCTGAGTAAC |
| | *Eco00* | AGACTGCGTACCAATTC |
| AFLP Amplification | *Mse*47 | GATGAGTCCTGAGTAAC AA |
| | *Mse*48 | GATGAGTCCTGAGTAAC AC |
| | *Mse*49 | GATGAGTCCTGAGTAAC AG |
| | *Mse*50 | GATGAGTCCTGAGTAAC AT |
| | *Mse*51 | GATGAGTCCTGAGTAAC CA |
| | *Mse*52 | GATGAGTCCTGAGTAAC CC |
| | *Mse*53 | GATGAGTCCTGAGTAAC CG |
| | *Mse*54 | GATGAGTCCTGAGTAAC CT |
| | *Mse*55 | GATGAGTCCTGAGTAAC GA |
| | *Eco*31 | 6-FAM-GACTGCGTACCAATTC AAA |

**Supplementary Table 2.** Chemico-physical composition of mineral water used as aqueous medium for *Lemna* clone growth.

| **Parameter** | **Value** |
|---|---|
| Temperature at source | 17.0 °C |
| Fixed residue at 180°C | 625 mg/L |
| Electrical conductivity at 20°C | 768 µS/cm |
| pH at source | 6.17 mg/L |
| Carbon dioxide $CO_2$ | 549 mg/L |
| Nitrites | < 0.002 mg/L |
| Nitrates | 38.8 mg/L |
| Silica | 64.0 mg/L |
| Hydrocarbon ions (bicarbonates) | 473 mg/L |
| Fluoro | 1.05 mg/L |
| Calcium | 97 mg/L |
| Magnesium | 24 mg/L |
| Potassium | 59 mg/L |
| Sodium | 45 mg/L |

**Supplementary Table 3.** Summary of the NewHybrids analysis for 5 random datasets consistent of 200 random AFLP loci each. Shown is the individual assignment to genomes (G = *L. gibba*, M = *L. minor*). All individuals were assigned with a high posterior probability (pp) > 0.999.

| | Expected ancestry proportions | | | | Everage | |
|---|---|---|---|---|---|---|
| **Genotype** | **GG** | **GM** | **MG** | **MM** | **pp** | **n** |
| *L. gibba* | 1 | 0 | 0 | 0 | 1.000 | 7 |
| *L. mediterranea* × *L. gibba* | 0.5 | 0.25 | 0.25 | 0 | 0.000 | 0 |
| *L. mediterranea* (F1) | 0 | 0.5 | 0.5 | 0 | 1.000 | 7 |
| *L. mediterranea* (F2) | 0.25 | 0.25 | 0.25 | 0.25 | 0.000 | 0 |
| *L. mediterranea* × *L. minor* | 0 | 0.25 | 0.25 | 0.5 | 0.000 | 0 |
| *L. minor* | 0 | 0 | 0 | 1 | 1.000 | 7 |

**Supplementary Table 4**. P value of ANOVA and Chi-squared Test for quantitative and qualitative traits respectively (*** for $p < 0.001$; ** for $p < 0.01$; * for $p < 0.05$).

| **Quantitative traits** | **p value** | |
|---|---|---|
| Frond length | 0.09801 | |
| Frond width | 3.746e-10 | *** |
| Frond length/width ratio | 0.0003767 | *** |
| Frond area | 8.027e-08 | *** |
| Vein number | < 2e-16 | *** |
| Root length | 0.008559 | ** |
| Distance frond base-root base | 0.3565 | |
| Aerenchyma % | 1.555e-11 | *** |
| Aerenchymatic cell length | 5.153e-11 | *** |
| **Qualitative traits** | **p value** | |
| Frond shape | 2.478e-05 | *** |
| Frond symmetry | 0.2013 | |
| Hyaline edge | 0.001734 | ** |
| Aerenchyma position | 2.2e-16 | *** |
| Aerenchyma reaching edge | 4.877e-06 | *** |
| Connection stipe presence | 0.0003443 | *** |
| Frond apex | 1.0000 | |
| Papules | 1.132e-06 | *** |

**Supplementary Table 5**. P values of Post-hoc for quantitative traits resulted significant from ANOVA test.

| **Frond width** | **GG** | **GGM** | **MG** |
|---|---|---|---|
| GGM | 0.461 | - | - |
| MG | 2.7e-05 | 1.0e-06 | - |
| MM | 3.3e-06 | 1.0e-06 | 0.013 |
| **Length/width frond ratio** | | | |
| GGM | 0.9893 | - | - |
| MG | 0.0274 | 0.0014 | - |
| MM | 0.0274 | 0.0018 | 0.7511 |
| **Frond area** | | | |
| GGM | 0.22867 | - | - |
| MG | 0.00014 | 7.3e-05 | - |
| MM | 7.4e-05 | 7.4e-05 | 0.08612 |
| **Aerenchyma %** | | | |
| GGM | 0.00018 | - | - |
| MG | 2.2e-08 | 0.00064 | - |
| MM | 8.1e-07 | 0.00011 | 0.01456 |
| **Longest aerenchyma cell** | | | |
| GGM | 3.2e-05 | - | - |
| MG | 9.8e-08 | 1.00000 | - |
| MM | 1.0e-06 | 0.00043 | 4.6e-05 |
| **Vein number** | | | |
| GGM | 1.00000 | | |
| MG | 0.00000 | 0.00000 | |
| MM | 0.00000 | 0.00000 | 6e-07 |
| **Root length** | | | |
| GGM | 0.84030 | - | - |
| MG | 0.00017 | 6.5e-05 | - |
| MM | 0.00922 | 0.00922 | 0.84030 |

**Supplementary File 1.** Nucleotide sequences gained by both the intronic regions of the β-tubulin locus *TUBB*1 and plastid markers (a*tp*F-*atp*H and *psb*K-*psb*I) for *L.* × *mediterranea* LM0027

>*TUBB*1-I-LM0027-subgenomeM
AGTGAGTTGATAGAGTGAGAAAACAGAGTATTATGATCTCGTTTAGTTTTACAATGATAGC
GATAAAAGTAATCTTGATCTATAACACTGATCCAAAAGAAATGGTTGACAAGATGAGACTC
CCAAAGGCGCATGGGTGGGATATTATGGAATAATGAGTGTGTGGTTATTTGTGTGAGAGAG
AGGGTGCATGTGATAGGGAGGGAGAGAATCCGCATGAGAGAGAGAGGGTGAGAGTATGT
GAACGTGTGTGAGAGAGAGGGAGAGGGAGGGAGGGGGGGGGAGAGAGAGAGAGAGA
GAGAGAGAATGTGGTGAGAGAGACTGTGAATGAGAGAGAGAGCAGAATTAAAAAGGGAA
AACAAACATAAAATGTCGAAGTACTATCTATAAACTCCAAAACGCCAATAAACTAACATA
AAACTGCTTGAGTGAGAACGAGTATAGATCTAAGTCGACTGAAAACCCTGGGAATCTAAC
ATAAA

> *TUBB* 1-I-LM0027-subgenomeG
TATACGATCTCATTCAGTTCTAAAATGATAGCGATCAAAGTAATCTTGATCTATAACACTG
ATTCAAGATAAATGGTTGACAAGAAGTGTGAGTGTTTGTGTGAGAGAGAATCCGCATGAG
AGAGAGGGAGAGACTACATGAACATGTGTGAGAGAGGGAGAGGAAAGTAGGGAAAGAGA
AAGAACACAGAGAGAAAGAGATAGAAAGAGAAAGAGTGAGAATGTGGGTGAGAGAGACT
GTGTATGAGAGAGACAAATGGTTGCCAAGATGAGAATCCCGATGGCAACATGGTTGACTA
GATGAGAGAGAGAAATGGAATATTGAGTGCGTGTGGGTGTTTGTGTTAGAGAGAGGGTGT
ATGTGAGAGGGAGAGGGAGAGAGAGAGAGAGAATGTGGGTGAGAGTGTGTGTATGAG
AGAGAGAGAGAGCAGAATTAAAAAGAAAAAAACATAAATTCTCGAAATATTATCTATAAA
GTCTAAAACGTCAATAAACTAACATAAAAATGCTCTAGTGAAAGCGAGTAAAGATCTAAG
TCGAATGAAAGCCATG

> *TUBB* 1-II-LM0027-subgenomeG
CCGACTATAAGAAATCAAACCAGAAACTTCTGTCAATTTTATGGTATTTGAATGAAACTTA
CGTTAACATGGGAATTAAGTAGAAAGAGCAAGCGGAAGAGAGAGAAATAATTGAACTCCT
CTCTCTCTCTCTCTCTAGGTGTATGTTCTTCTCAGATCATCAACTATAATAAAAAAATGTGA
TGTTTTTGGAAAATAGATAAAATTTACGGATCTTACAGCTTGG

> *TUBB* 1-II-LM0027-subgenomeM
GTTCTGTCAATTTTATGATATTTAAATGAAACTTAAATCAACATGTGAAAACACTTAGATCC
CACTTGGCAATGTTAGTAAAGTTTCATGGGAGTTCTAGAGAGGGAGAGGGGGGGTTGAA
CTCGTAAGAAACAATCTCCTCTCTCTTTTTCTCTCTTCACATGGATAATCATTCAGTCTCAG
ATAATATCTTACTATCTAACATGGTCACATTACCAACATCTTCATCTTCATCATGCATCCTA
ACCAAAAACACAAACGATGATAAAACATGTATAATCATCATGGCTATTTAAGTAAAGCTTC

AAGGGAATTCTAGAGAGCGAAAGAGAGAGAGTTGAACTCCCAACGGAATTCTACGTTTGT
TCAACTTCCTCCATGTCAAATGAAATCTAATGTGCCAACGTCCAAGAGCATGGGAATTCTA
GAGAGAGCAAGAGAAAGAGAGAAAAGAATTGAACTTCTCTCTCTCTCCCCCTCTCCAGGC
ATATGTTCATCCAATCTCAGATCGTCAACTA

>LM0027-atpF-H

ATTCTAGAATAAAAAAGTACGTAATAGACTTTTTGACTTAGACTTGCTTTTTGCTTCTTCGA
ATTATATCAACATTGTACTCTAACAATTACTTATTCGTTGAGAGAATACCTCCGGGAAGGA
CTGATTTTAGGATTAGTAATTAGCAGATCCTCTCGCTTTCTTCCTTCCCGTTTTCAGTTCTTA
GTATAATGTAATGCAAAACTTTTTTGAGTATGCGTTGTAACGCAATAAACAAGGTATTTAC
CAATTGACAAAATAGCCAGGACCTAACCCAATAAGTATGTTCTTGTAATTGTAAACTTTAA
TTAGAATTAAATAATAAATAAATAAAGTTCTCAATTAAATTAATTAATTTAATCTATTCCAT
TTTTAAATCCCATAAAAAAAAAAAAAGAAATCAAACAAAGGGGGCGAAGTAATACAAAAG
GAACTCTGTTCTTTTTTAGTCCTATCTATAAGAGGAGAGTATATGCAAAATGTAACTGATTC
TTTTGTTTCCTTGGGCCACTGGCCATCCGCCGGCGGTT

>LM0027-psbK-I

TGAGTTTTCGATAAAGTTATTAAAACTCTACTGAAAAAATTCATGATTTATTTGATAAAAA
AGATTCTAATAAAAATTGATAACGTAGTAGCAATCTTAGTTTATACATCCTCATAAAAAAT
ATTTGAATTCTTGTATATTGGATAAAAAGAGCGATAAGTTTGGATCAGTCCATTTCGCCGTT
CTGGCCGCTCTTCCAGTGAGGAAGTACTTTATTTATTTATTAGCTTTTGTTTTACACAATACT
TTATTGTTAATATTAGAGTTAATATTAGAATAACCTTTTTGGTAACGAACAAGTCATAATCT
TAATTTAAAATGCATTCATGAGTTTGAAAATTCAGTTTTTGTAGAAAAAAACACTAAACTT
AATACTATACTTAATAAAATAAAGAATTGTTCTTTATTTTTTCATAGTTTTTTTTCTTGGCAT
GCCCAAATAATACATGTGTTACATAACTCAAATGGATAATCTATTCCCTTTTACCCCAAAA
ATGATCCTATCTTGGAG
